# Retinal Calcium Waves Coordinate Uniform Tissue Patterning of the *Drosophila* Eye

**DOI:** 10.1101/2025.06.02.657513

**Authors:** Ben Jiwon Choi, Yen-Chung Chen, Claude Desplan

## Abstract

Optimal neural processing relies on precise tissue patterning across diverse cell types. Here, we show that spontaneous calcium waves arise among non-neuronal support cells in the developing *Drosophila* eye to drive retinal morphogenesis. These waves are initiated by Cad96Ca receptor tyrosine kinase signaling, triggering PLCγ-mediated calcium release from the endoplasmic reticulum. A cell-type-specific ‘Innexin code’ coordinates wave propagation through a defined gap junction network among non-neuronal retinal cells, excluding photoreceptors. Wave intensity scales with ommatidial size, triggering stronger Myosin II-driven apical contraction at interommatidial boundaries in larger ommatidia. This size-dependent mechanism compensates for early boundary irregularities, ensuring uniform ommatidial packing critical for precise optical architecture. Our findings reveal how synchronized calcium signaling among non-neuronal cells orchestrates tissue patterning in the developing nervous system.

## INTRODUCTION

Precise tissue patterning is crucial for optimal biological function (*1–3*). Sensory systems exemplify this principle with their arrangement of functional modules that is essential for processing sensory input (*4, 5*). To organize sensory modules, non-neuronal cells like glia and epithelia must be precisely patterned and coordinated with sensory neurons (*6–8*).

Synchronized calcium activity in neuronal cells during development is crucial for the patterning of neuronal circuits (*9–12*). In the mammalian visual system, retinal waves guide wiring of visual pathways within the retina and in higher brain areas (*13–15*). While also affecting non-neuronal cells like Müller glia and pigment epithelium, their role in organizing nervous system architecture remains unclear (*16–18*). Beyond the nervous system, synchronized calcium activity is critical for morphogenesis in other tissues (*19–22*), for instance coordinating myoblast fusion into muscles (*23*). This suggests it may also guide structural patterning of non-neuronal cells in the nervous system.

In contrast to synchronized calcium activity mediated by synaptic transmission between neurons, this activity in non-neuronal cells is primarily mediated by gap junction channels (*19–21*) that form multicellular networks (*24, 25*). Beyond their role in nervous system development, gap junction networks are crucial for various biological processes, including tissue regeneration, cancer metastasis, and aging (*26–28*). However, technical difficulties have hindered a comprehensive understanding of their roles. Challenges include complex live imaging and difficulty manipulating gap junctions due to subunit diversity and essential roles in cellular physiology (*24, 29*).

The *Drosophila* retina offers an excellent system for investigating the role of intercellular calcium activity among non-neuronal cells (*30–32*). Its simplicity, stereotyped organization, and genetic accessibility enable precise live imaging and manipulation. It has recently been used to identify the role of calcium waves during late retinal development (*33*).

The fly compound eye has ∼800 unit-eyes (ommatidia), each of which houses eight photoreceptor neurons (**PRs**) (*34, 35*). These units are surrounded by specialized non-neuronal retinal support cells that exhibit features of glial and epithelial cells (**Fig. 2A-H**) (*36, 37*). They include four cone cells (**CCs**), two primary pigment cells (**1PCs**), and a cluster of twelve interommatidial cells (**IOCs**) that are shared among six adjacent ommatidia (*35, 37*): six secondary pigment cells (**2PCs**), three tertiary pigment cells (**3PCs**), and three bristle cells (**BCs**).

Both neuronal and non-neuronal cells show regional patterning: the dorsal rim detects polarized light (*34*), and dorsal R7 photoreceptors co-express two UV opsins (*38*). The ventral retina features structurally larger ommatidia compared to the dorsal side (*39, 40*). This regional patterning is evolutionarily conserved across fly species and is linked to optimized visual detection of low intensity ground features (*39*). The optic lobe circuitry also exhibits regional polarity, with distinct synaptic densities and neuronal subtypes between the dorsal and ventral sides (*41, 42*). These findings indicate robust regional patterning of neuronal and non-neuronal populations critical for visual processing (*43*).

In this study, we monitored live retinal calcium dynamics throughout the pupal stages. We identified robust early retinal waves of synchronized calcium activity in retinal support cells, just prior to synaptogenesis of PRs to their optic lobe targets. We investigated the molecular mechanisms and the developmental role of these waves. Calcium release from the endoplasmic reticulum via IP_3_ receptor (IP_3_R) is initiated by the receptor tyrosine kinase (RTK) Cad96Ca and phospholipase Cγ. It propagates to adjacent cells through gap junctions, with unique Innexin subunit combinations involved in distinct cell types. Retinal waves drive calcium-dependent apical contraction of interommatidial boundaries through cell morphogenesis via the Myosin II pathway. Wave activity is more frequent in the ventral region of the eye, where larger ommatidia generate stronger calcium signals that promote more robust contractions. This mechanism compensates for initial boundary irregularities, ensuring uniform interommatidial boundaries across the retina, which is critical for establishing consistent lens architecture. Therefore, we reveal the molecular and cellular mechanisms by which synchronized calcium activity in retinal support cells coordinate the overall architecture of the eye.

## RESULTS

### Early retinal calcium waves in the developing retina

We utilized a dry-lens imaging platform to monitor calcium dynamics in the developing retina of intact animals **(Fig. 1A**). Compared to the immersion-lens protocol (*44, 45*), this approach eliminates the need for coverslips, enabling simpler and contact-free imaging of intact animals **(Fig. 1A, B**). We monitored calcium dynamics in the developing retina from ∼10 hours after pupal formation (**10 hAPF**) until eclosion (∼100 hAPF) **(Fig. 1C, S1A-K, Movie 1**). We expressed the genetically encoded calcium indicator GCaMP6s in all retinal cells (*46, 47*).

**Fig. 1.**
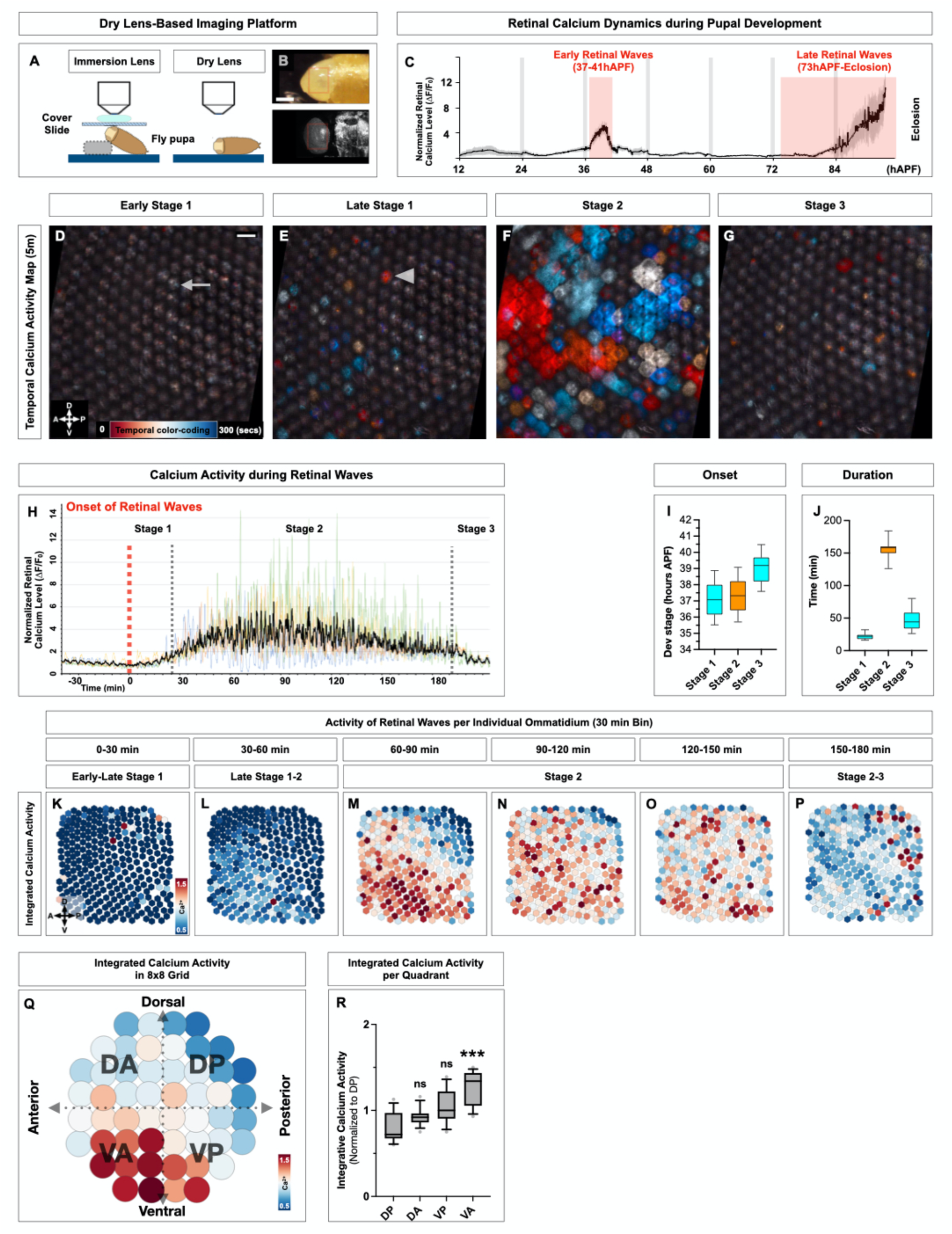
Retinal calcium wave dynamics during pupal eye development (A-R). **1A.** Schematic of imaging platform using a dry-lens setup for intact pupa, shown in comparison with a conventional immersion-lens setup. **1B.** Representative images of sample preparation (top) and retinal calcium imaging (bottom). The red box marks the retinal area analyzed in **(D–G)**. Scale bar: 500 μm. **1C.** Average calcium dynamics across pupal retinal development. Imaging was performed every 2 min, and n ≥ 5 retinas were analyzed for each time window. Details are provided in **Fig. S1A–K.** See also **Movie 1**. **1D–G.** Retinal calcium wave activity shown as color-coded 5-min temporal activity maps (T-stacks) for early stage 1 **(D)**, late stage 1 **(E)**, stage 2 **(F)**, and stage 3 **(G)**. The arrow in **(D)** indicates a single cone cell firing, and the arrowhead in **(E)** indicates full activation of an ommatidium. Images are from the same retina, imaged at 3-s intervals. (A = anterior, P = posterior, D = dorsal, V = ventral). Scale bar: 20 μm **(D–G)**. See also **Movie 2**. **1H.** Individual calcium traces corresponding to **(D–G)**. Four retinas are shown in different colors, and the black trace represents the averaged activity. **1I.** Quantification of retinal wave onset timing (n = 19). Error bars: SD. **1J.** Quantification of retinal wave duration (n = 19). Error bars: SD. **1K–P.** Ommatidial-scale integrative calcium activity between 37–41 hAPF. Each bin corresponds to a 30-min period, and each hexagon represents a single ommatidium. Signal intensity for each ommatidium is color-coded. Data are from the same retina shown in **(D–G)**. **1Q.** Grid-based map of integrative calcium activity. Each circle represents a group of ommatidia arranged in an 8 × 8 grid along dorsoventral (**DV**) and anteroposterior (**AP**) axes. Average intensity of integrative calcium activity per grid point is color-coded. n = 6 retinas. **1R.** Quantification of quadrant-specific integrative activity shown in **(Q)**. n ≥ 432 ommatidia from 6 retinas. (ns: p ≥ 0.05, *p < 0.05, ***p < 0.001). Error bars: SD.

During our observations, two specific developmental windows displayed robust synchronized calcium activity across the whole retina **(Fig. 1C, S1I, K).** Early retinal waves, hereafter referred to as “retinal waves”, emerged at around 37 hAPF and lasted roughly 3 hours **(Fig. 1C, 1I, Movie 1H)**. Previously described and analyzed later-stage calcium waves were observed starting from around 73 hAPF and continued until eclosion **(Fig. 1C, S1K, Movie 1I**) (*33*).

We focused on the early retinal waves with fine temporal resolution to determine their overall properties and function **(Fig. 1D-J, Movie 2**). During early pupal development before the onset of retinal waves (37 hAPF), isolated calcium transients were only rarely observed in **PRs (Fig. 1H arrow**). Retinal waves began as calcium signals in a single retinal cone cell, appearing at 37.1 ± 0.91 hAPF (SD, n=19), initiating what we define as early stage 1 **(Fig. 1D arrow, H-J, Movie 2A, Fig. 2**). The signals then spread to adjacent cells within the same ommatidium during late stage 1 **(Fig. 1E, Movie 2B**). These stage 1 waves lasted for 22±4.4 minutes **(Fig. 1J**). Then, the waves started spreading to adjacent ommatidia, which marked the onset of stage 2 **(Fig. 1F, H, Movie 2C**). This phase was characterized by intense retinal calcium activity that persisted for 156±14 minutes **(Fig. 1H, J**). Subsequently, at 39.1 ± 0.86 hAPF, the waves became again restricted to a single ommatidium for 46 ±15 minutes (stage 3) **(Fig. 1G, H-J, Movie 2D**). Later, most calcium activity faded, leaving sporadic calcium transients, mainly in **PRs**, which continued until the onset of the late retinal waves at 73.4 ± 1.26 hAPF (SD, n=7) **(Fig. S1K)** (*33*). The onset and duration of retinal waves were consistent between animals **(Fig. 1H-J**), occurring irrespective of the genetic drivers, calcium sensors, and excitation laser used **(Fig. S1M-O)**. When we simultaneously imaged both eyes, the difference in the onset of the waves between eyes was only 2.4±1.7 minutes (SD, n=5), suggesting that the initiation of retinal waves is a bilaterally synchronized phenomenon **(Fig. S1L, Movie S1**).

**Fig. 2.**
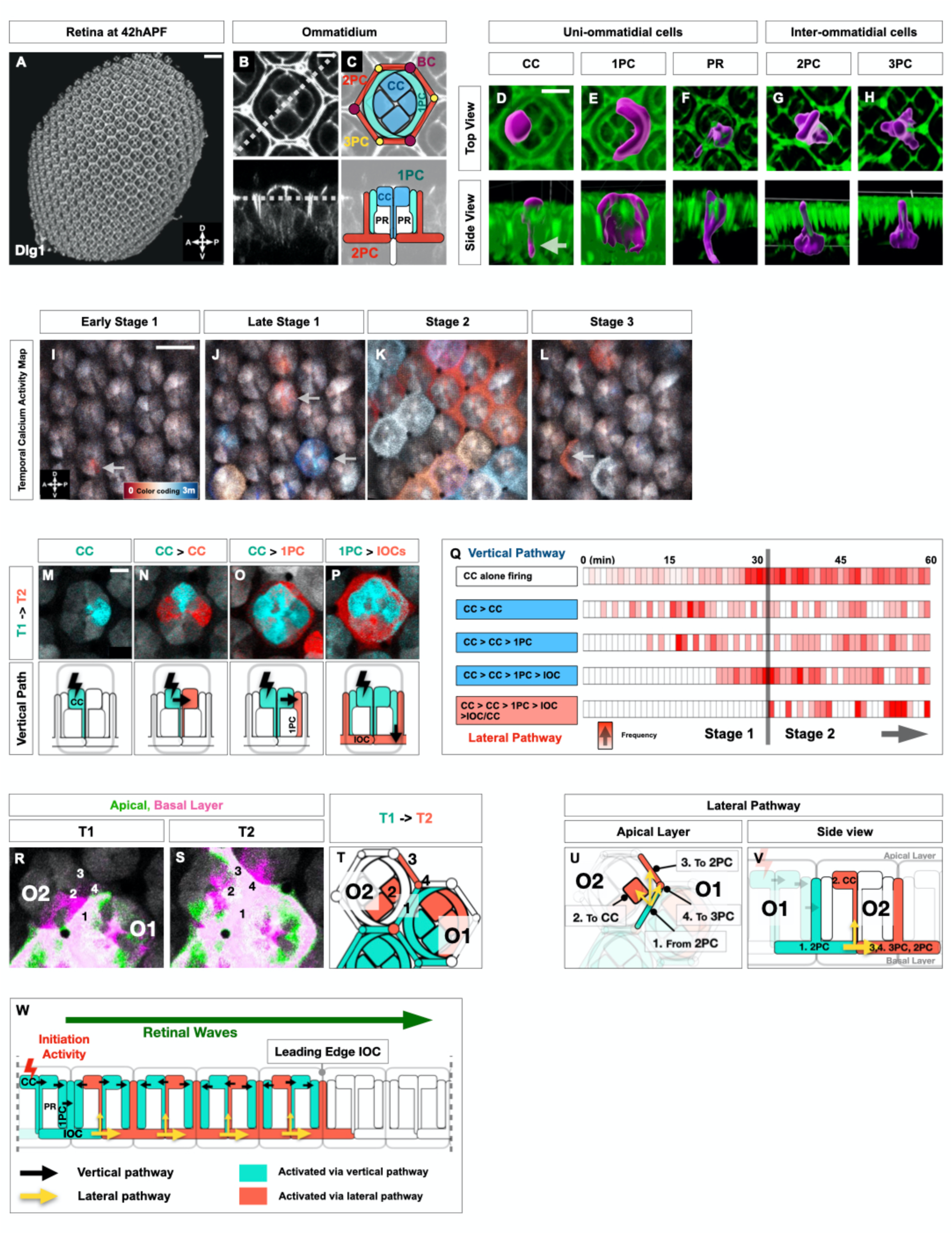
Single-cell resolution of retinal waves reveals stereotyped initiation and propagation patterns (A-W). **2A–C.** Retina overview **(A)**, apical side **(B top)** and cross-section **(B bottom)** of a single ommatidium immunostained with Dlg1. The dotted line in **(B top)** indicates the cutting plane for **(B bottom)** and vice versa. **(C)** Diagram of retinal cell types: cone cell (CC), primary pigment cell (1PC), secondary pigment cell (2PC), tertiary pigment cell (3PC), bristle cell (BC), photoreceptor (PR). Scale bars: 20 μm **(A)**, 5 μm **(B, C)**. **2D–H.** Three-dimensional reconstructions of retinal cell morphology in *longGMR-GAL4>UAS-MCFO* retina at 40 hAPF. Top **(D–H tops)** and side views **(D–H bottoms)** show single cell morphology. Magenta marks single-cell labeling, green marks Dlg1. Arrow in **(D bottom)** highlights the leg-like structure of CC contacting IOCs. Scale bar: 10 μm **(D–H)**. **2I–L.** Retinal calcium wave activity at single-cell resolution displayed as color-coded 3-min temporal activity maps for early stage 1 **(I)**, late stage 1 **(J)**, stage 2 **(K)**, and stage 3 **(L)**. The arrow in **(I)** indicates a single cone cell firing, arrows in **(J)** indicate vertical waves, and the arrow in **(L)** indicates a primary pigment cell firing. All images came from the same retina. Scale bar: 10 μm **(I–L)**. See also **Movie 3**. **2M–P.** Vertical pathway patterns including single CC firing **(M)**, CC–CC propagation **(N)**, CC– 1PC propagation **(O)**, and CC–IOC propagation **(P)**. Time points are shown at T1 (cyan) and T2 (red, 4 s later). Bottom panels show schematics. Scale bar: 5 μm **(M–P)**. See also **Fig. S2A, C-E**, **Movie 4**. **2Q.** Event log of retinal waves from stage 1 through early stage 2 in a single retina. Each bin represents a 1-min interval, and event frequency is color-coded. Blue box marks the vertical pathway, and red box marks the lateral pathway. **2R–T.** Lateral pathway wave patterns. **(R, S)** show T-series with apical (green) and basal (magenta) layers, where T1 and T2 are separated by 4 s. Cells: initial 2PC (1), propagated CC (2), 2PC (3), 3PC (4). **(T)** shows schematic representation of calcium signaling at T1 (cyan) and T2 (red). Scale bar: 5 μm **(R, S)**. See also **Fig. S2F**, **Movie 5**. **2U, V.** Diagrams of lateral pathway wave patterns as in **(R–T)**. **2W.** Diagram of retinal wave propagation along vertical and lateral pathways across multiple ommatidia.

We examined regional patterns of retinal waves by quantifying the amount of synchronized calcium activity occurring across multiple cells within each ommatidium. During late stage 1, calcium activity first emerged in isolated ommatidia, predominantly in the ventral region **(Fig. 1L, M, S1P, Q**). As the waves progressed to stage 2, they spread across neighboring ommatidia to encompass other retinal regions **(Fig. 1M, N**). To analyze wave activity across the population, we integrated data from multiple retinas by pooling them into an 8×8 grid template **(Fig. 1Q**). This spatial analysis over the entire wave period revealed a distinct regional pattern, with increased wave activity in the ventral-anterior (**VA**) region, further explored in **Fig. 5**. **(Fig. 1Q, R**).

### Single-cell resolution of retinal waves reveals stereotypical patterns of initiation and propagation

To examine retinal wave initiation and propagation at the single-cell level, we used multi-color Flp-out labeling at 40 hAPF to map individual cell configurations **(Fig. 2A-H**) (*48*). Within a single ommatidium, all retinal cell types are in direct contact with each other **(Fig. 2C**), consistent with electron microscopy studies (*35*), providing a structural basis for single-cell tracing of retinal waves.

We performed high-resolution calcium imaging to detect single-cell calcium activity **(Fig. 2I-L, Movie 3**). Initial calcium activity began from single cone cell (**CC**) **(Fig. 2I arrow, 2M**), with all four cone cells capable of initiating with a slight bias toward dorsal-facing cone cells (**Fig. S2B**). It then propagated stereotypically to neighboring cells **(Fig. 2M-P**). During Stage 1, the waves began in one cone cell, spread to adjacent cone cells (**Fig. 2N**), then moved to primary pigment cells (**1PCs**) (**Fig. 2O**) before reaching interommatidial cells (**IOCs**) (**Fig.2P**), thus propagating within a single ommatidium (**CC>CC>1PC>IOCs**) **(Fig. 2M-Q, S2A, C-E, Movie 4**). Calcium thus spread to all non-neuronal cell types in an ommatidium without involving photoreceptor neurons (**PRs**). These stereotypic propagation patterns progressed from the apical to the basal layer **(Fig. 2M-P**), defining **CC>CC>1PC>IOCs** as a ‘vertical pathway’ (**Fig. 2W**). Although all retinal cells within a single ommatidium are in physical contact **(Fig. 2C**), no alternate wave routes were observed, such as **1PC>CC** or **CC>IOC** bypassing **1PC**.

Stage 2 calcium activity spread laterally to neighboring ommatidia, mediated by interommatidial cells as a ‘lateral pathway’ **(Fig. 2K, R-W**). Two adjacent ommatidia O1 and O2 share interommatidial cells, allowing interommatidial cell in one ommatidium (**O1**) to directly contact interommatidial cells and cone cells in **O2 (Fig. 2R-V**). During wave propagation from **O1** to **O2 (Fig. 2R, S**), the calcium signal spread from this initial interommatidial cell (**2PC** labeled ‘**1’**) to cone cells (‘**2**’) and interommatidial cells (‘**3**’ and ‘**4**’) through basal layer connections **(Fig. 2U, V**). The lateral propagation moved across successive ommatidia, with interommatidial cell–to–cell signaling forming the leading edge of the wave **(Fig. 2W, S2F, Movie 5**). Meanwhile, interommatidial cells also activated cone cells along the wave’s path, activating all other cells and initiating a vertical pathway within each ommatidium. This coordination activated all support cells along the wave path, establishing robust, region-specific calcium signaling **(Fig. 2W**).

Vertical and lateral waves then stopped, leaving sporadic calcium activity in cone cells and primary pigment cells that defined stage 3 **(Fig. 2L, Movie 3D**). We never observed calcium activity or propagation to **PRs**: GCaMP6s driven by a **PR**-specific GAL4 driver (Chaoptin-GAL4) did not show any calcium activity in **PRs** during the waves **(Fig. S3A-C, Movie S2**): Retinal waves are entirely driven by non-neuronal retinal cells, similarly to late retinal waves (*33*).

In summary, retinal waves do not diffuse randomly despite full cell interconnections; they initiate in cone cells and follow organized vertical and lateral pathways **(Fig. 2W**), suggesting an underlying dedicated cellular communication network.

### Cad96Ca-PLCγ-IP_3_R signaling triggers calcium release from the ER

In mammals, the endoplasmic reticulum (**ER**) is the primary source of calcium for the waves in glia and epithelia (*21, 49*), suggesting a similar role in *Drosophila*. To simultaneously measure calcium activity in both the ER and the cytoplasm, we used two calcium indicators with different colors: ER-targeted GCaMP (green), and cytosolic jRGECO1a (red) **(Fig. 3A-D, Movie 6**) (*50–52*). During the waves, cytoplasmic calcium rose as ER calcium signal fell across all retinal supporting cells **(Fig. 3A-D**), indicating active ER calcium release.

**Fig. 3.**
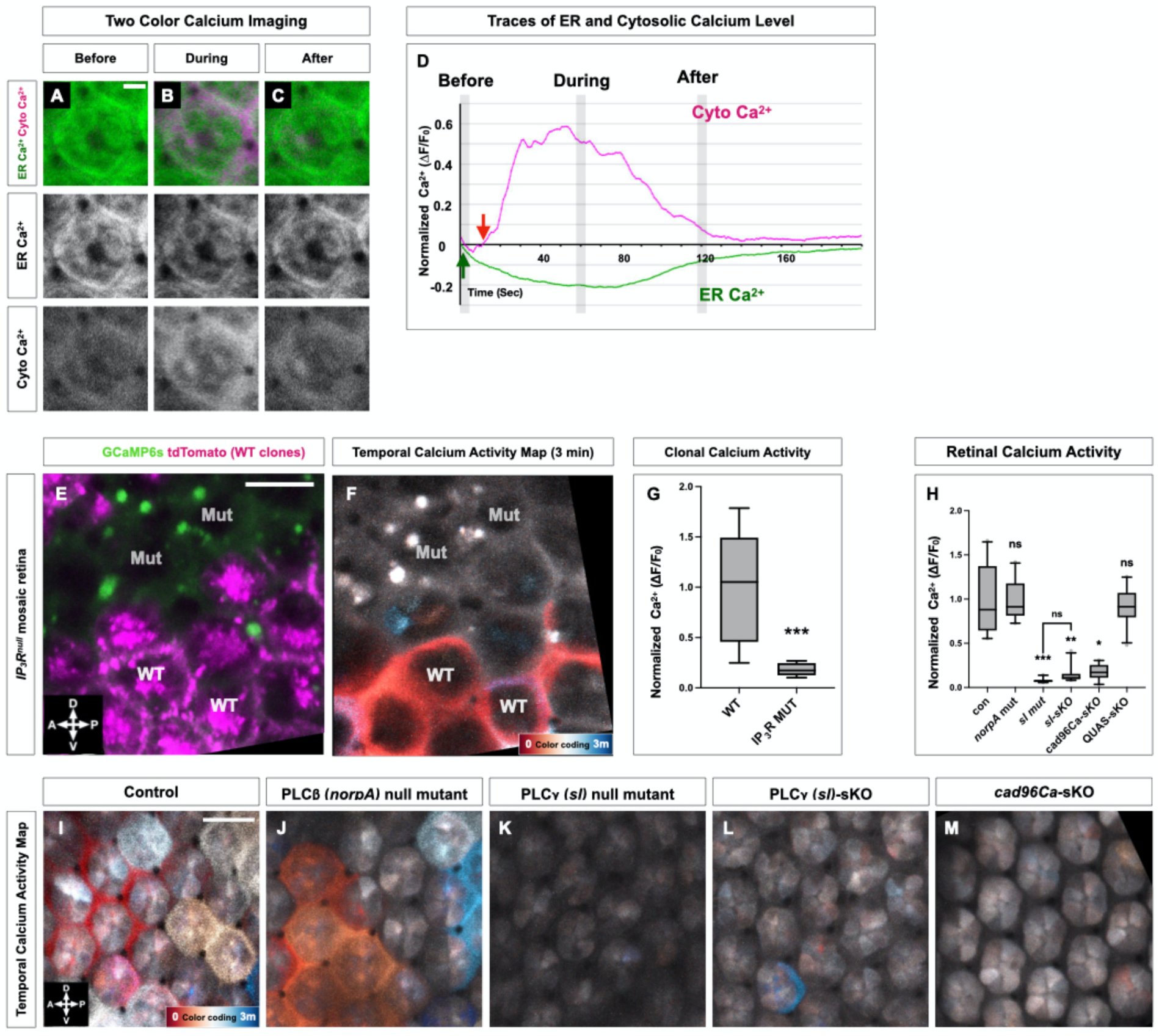
Cad96Ca-PLCγ-IP₃R signaling triggers ER calcium release, initiating retinal waves (A–M). **3A–C.** Dual-color calcium imaging before, during, and after retinal waves, selected at 60-s intervals. Red: cytosolic jRGECO1a; green: ER-targeted GCaMP6. Imaging was performed at 1-s intervals at the retinal center during stage 2. Scale bar: 5 μm. See also **Movie 6**. **3D.** Calcium traces during retinal waves. Shaded regions correspond to snapshots in **(A–C)**. Arrows indicate wave initiation in each channel. **3E, F.** IP₃R mutant mosaics. **(E)** Wild-type (magenta) vs mutant (non-labeled) clones. **(F)** Three-min temporal activity maps using *longGMR-GAL4>UAS-GCaMP8s* in mosaic retina. Scale bar: 10 μm **(E, F)**. See also **Movie 7**. **3G.** Quantification of calcium activity in wild-type vs mutant ommatidia from IP₃R mosaics. n ≥ 8 ommatidia from 3 retinas. ***p < 0.001. Error bars: SD. **3H.** Quantification of retinal wave activity from **(I–M)**. n ≥ 10 retinas. ns: p ≥ 0.05, ***p < 0.001. Error bars: SD. **3I–M.** Retinal calcium waves displayed as color-coded 3-min temporal activity maps in control **(I)**, *norpA* **(J)**, *sl* **(K)**, *sl-sKO* **(L)**, and *cad96Ca-sKO* **(M)**. sKO = CRISPR-mediated somatic knockout. Scale bar: 10 μm **(I–M)**. See also **Movie 8**.

We analyzed the role of the inositol trisphosphate receptor (**IP_3_R**), a crucial component of calcium wave dynamics (*49*). We generated retinal FLP/FRT mosaic clones, marking wildtype clones with RFP and leaving IP₃R mutant clones unmarked **(Fig. 3E**) (*33, 53*). Retinal waves failed to initiate or propagate in *IP_3_R* mutant clones, even at the borders of the clones **(Fig. 3F, G, Movie 7**), indicating that IP_3_R-mediated ER calcium release is cell-autonomously required for both the triggering but also the propagation of retinal calcium waves, likely through the sustained release of calcium from the ER in response to stimulation of IP_3_R.

We then investigated Phospholipase C (**PLC**) that generates **IP_3_** from phosphatidylinositol 4,5-bisphosphate (PIP_2_) (*54*). Among the five *Drosophila* PLCs, only PLCβ (*norpA*) and PLCγ (*sl*) are expressed in the retina (*55, 56*). *norpA* null mutants exhibited normal wave activity, but no waves were observed in *sl* null mutants **(Fig. 3J, K, H, Movie 8**). Although *sl* null mutants are viable, they exhibit impaired retinal growth **(Fig. S3E, H**) (*56*).

We therefore used longGMR-Cas9 to generate *sl* CRISPR-mediated somatic knockout (**sKO**) in mid-pupal retinas (*57*), bypassing early retinal development. *sl* somatic knockout retinas exhibited slightly reduced growth **(Fig. S3F, H**) but showed a 90% decrease in waves **(Fig. 3L, H, Movie 8D**), confirming the specific role of PLCγ in wave activity.

RTKs are upstream regulators of PLCγ signaling (*58*). We screened fifteen RTKs expressed in the developing retina (*59*) via somatic knockout, RNAi, or dominant negative transgenes (**Table S1**). Only *cad96Ca* (*stit*) was necessary for retinal waves initiation without altering gross retinal morphology **(Fig. 3M, H, S3G, H, Table S1**). *cad96Ca* is a cadherin-like RTK involved in wound healing (*60, 61*) with no known role in retinal development. *cad96Ca*-GAL4-driven GFP revealed expression in all support cells, but not **PRs (Fig. S3I-L**). These results suggest that Cad96Ca activates the PLCγ-IP_3_R pathway, inducing ER-release calcium activity during retinal waves.

### Retinal waves propagate through gap junctions mediated by cell-specific combinations of Innexins

Gap junction mediate intercellular calcium wave propagation (*21, 62, 63*). In *Drosophila*, eight *innexins* (***inx***) encode subunits that assemble into octameric hemi-channels forming gap junction channels at cell junctions (*64–66*), with some expressed in the developing retina (*66, 67*). Among them, our immunostaining showed Inx1 (*ogre*), Inx2, and Inx3 at non-neuronal retinal cell junctions at 40 hAPF **(Fig. 4A-I, S4A-F**) (*66–68*). They were localized in puncta that represent gap junction plaques, specialized clusters of gap junction channels, adjacent to the junctional marker Disc large1 (Dlg1) **(Fig. 4A-E**) (*69, 70*). Super-resolution STED imaging confirmed Inx2 and Inx3 strictly confined to gap junction plaques **(Fig. 4A-D**), whereas Inx1, though mainly at gap junction plaques, was also seen elsewhere in the cell **(Fig. 4E, Inx1 row**). In contrast, we did not detect Inx4, 5, 6, 7, or 8 at retinal cell junctions **(Fig. S4G-K**). Inx8 (*shakB*) was only detected at 48 hAPF, after the waves, and only in synapses within the optic lobe **(Fig. S4L-N**) (*66, 67*). Retinal waves remained unchanged in *inx8* mutants **(Fig. S4O-Q, Movie S3**). Given that Inx2 and Inx3 were absent from gap junction plaques in **PR** somas (see below, **Fig. S4D, E, PR layer rows**), this suggests that neither Inx8 nor **PRs** are involved in retinal waves.

**Fig. 4.**
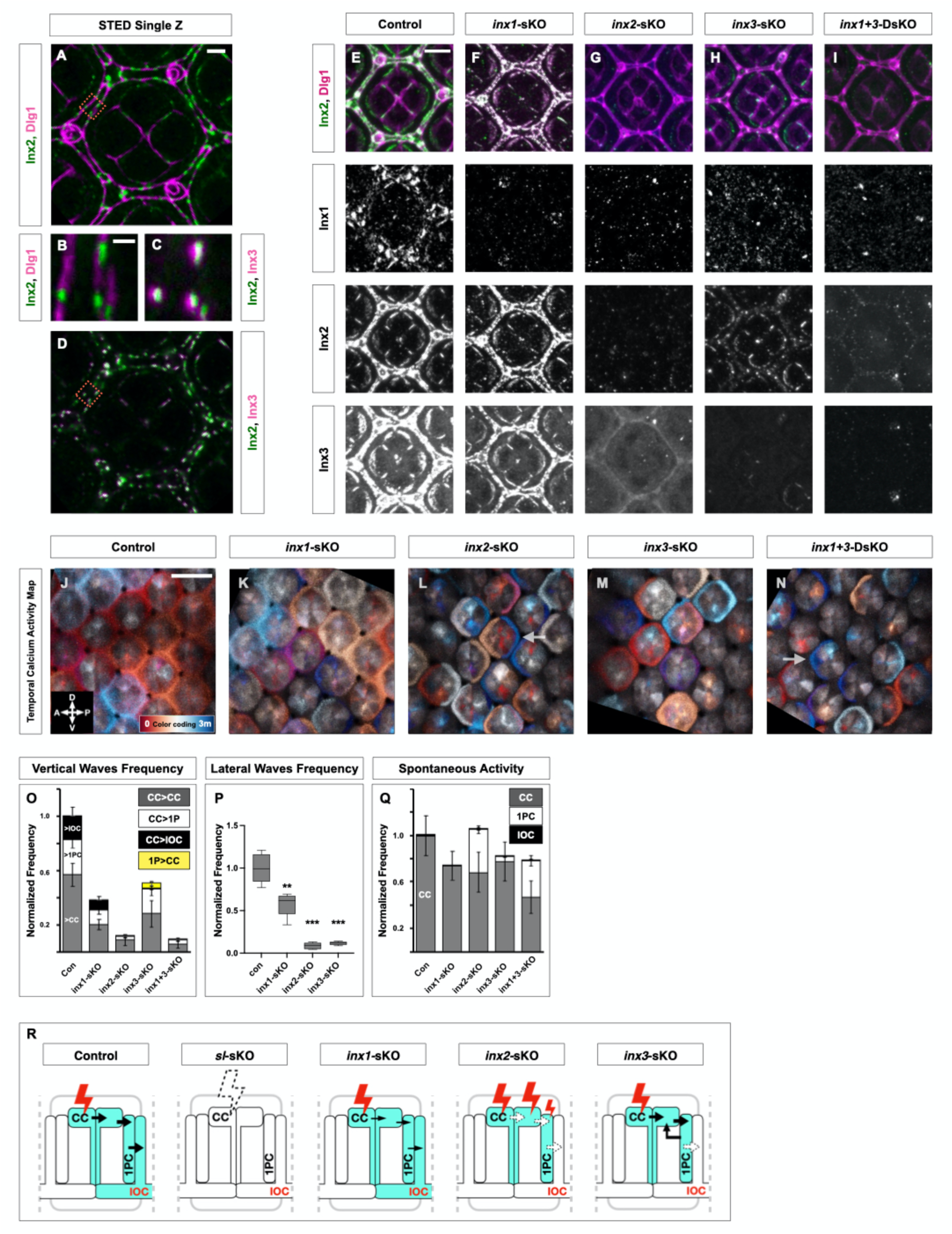
Retinal waves propagate through gap junctions using cell-type specific combinations of Innexins (A–R). **4A–D.** STED imaging of a single ommatidium at 40 hAPF stained for Dlg1 and Inx2 **(A, B)** or Inx2 and Inx3 **(C, D)**. Insets from (**A**) and (**D**) show the red-box regions in (**B**) and (**C**), respectively. Scale bars: 2 μm **(A, D)**, 0.5 μm **(B, C)**. **4E–I.** Immunostaining for Inx1, Inx2, Inx3 at 40 hAPF in control **(E)**, *inx1-sKO* **(F)**, *inx2-sKO* **(G)**, *inx3-sKO* **(H)**, and *inx1+3-double sKO* **(I)**. Green marks Innexins, and magenta marks Dlg1. Images are Z-stacks of the apical retinal layer. Scale bar: 5 μm **(E–I)**. **4J–N.** Retinal calcium wave activity as 3-min temporal activity maps in control **(J)**, *inx1-sKO* **(K)**, *inx2-sKO* **(L)**, *inx3-sKO* **(M)**, *inx1+3-DsKO* **(N)** during stage 2. Arrows in **(L, N)** indicate unsynchronized cone and pigment cell firing. Scale bar: 10 μm **(J–N)**. See also **Movie 9**. **4O.** Quantification of vertical pathway frequency in *inx* mutants during stage 2. n ≥ 5 retinas. Error bars: SD. Detailed statistics in **Fig. S4Z**. **4P.** Quantification of lateral pathway frequency in *inx* mutants during stage 2. n ≥ 5 retinas. ns: p ≥ 0.05, **p < 0.01, ***p < 0.001. Error bars: SD. **4Q.** Quantification of spontaneous cell-autonomous activity by cell type in *inx* mutants during stage 1. n ≥ 5 retinas. Error bars: SD. Detailed statistics in **Fig. S4Z**. **4R.** Diagram illustrating vertical propagation pathway in *inx* mutants.

To selectively inhibit *innexin* expression at mid-pupal stages, we generated somatic knockouts using longGMR-Cas9. Immunostaining confirmed the near-complete loss of the targeted *innexins* after somatic knockout **(Fig. 4F–H, Inx1–3 rows, S4W-Y**) in 40 hAPF retinas. Somatic knockout of *inx1, inx2, or inx3* did not alter cone cell activation **(Fig. 4Q, S4Z**), but each disrupted stereotyped steps of wave propagation **(Fig. 4J-P, R, S4Z, Movie 9**):

– Somatic knockout of *inx2* drastically disrupted the propagation of all waves **(Fig. 4L, O-R, S4Z**).
– Somatic knockout of *inx*1 only exhibited a slight decrease of the vertical and lateral wave pathways (**Fig. 4K, O-R, S4Z**).
– Somatic knockout of *inx3* retinas retained the initial vertical propagation from cone cells to primary pigment cells, but the propagation from primary pigment cells to interommatidial cells was severely disrupted (**Fig. 4M, O, R, S4Z**), resulting in a substantial reduction of the lateral pathway (**Fig. 4P**). Somatic knockout of *inx3* also exhibited reverse propagation from primary pigment cell to cone cell **(Fig. 4O**), which was never observed in wild type.
– Since somatic knockout of *inx1* and *inx3* each caused partial wave disruptions (**Fig. 4K, M, O**), we examined their double somatic knockout (***inx1+3*-DsKO**) **(Fig. 4I**) which led to complete disruption of wave propagation similar to somatic knockout of *inx2* (**Fig. 4N-Q, S4Z**).
– Somatic knockout of *inx2*, *inx3*, and the double somatic knockout of *inx1* and *inx3* caused little or no decrease in spontaneous firing in cone cells between 37 hAPF and 40 hAPF (**Fig. 4Q, S4Z**). However, infrequent ectopic firing was also observed in primary pigment cells and interommatidial cells (**Fig. 4Q**), suggesting that the waves normally suppress ectopic activity in non-cone cells.

To elucidate the distinct roles of each Innexin, we focused on the formation of gap junction plaques **(Fig. 4A-D**) that are critical for organizing gap junction channels (*69, 71*). Gap junction plaques disappeared in *inx2* somatic knockout retinas, leading to dispersed staining of Inx1 and Inx3 outside of cell junctions (**Fig. 4G, S4W-Y**) showing that Inx2 is important for their junctional localization (*68*). However, gap junction plaques persisted in *inx1* somatic knockout retinas but were slightly reduced in size based on Inx2 and Inx3 staining (**Fig. 4F, S4W-Y**). In *inx3* somatic knockout retinas, gap junction plaques were specifically lost in interommatidial cells, consistent with impaired lateral wave propagation, whereas smaller gap junction plaques remained in other cell types (**Fig. 4H, S4W-Y**). In the double somatic knockout retina of *inx1* and *inx3*, gap junction plaques were absent at all cell junctions, similar to *inx2* somatic knockout (**Fig. 4I, S4W-Y**). As expected, whenever gap junction plaques disappeared upon somatic knockout of a given *innexin*, the non-targeted Innexins were still detectable but were dispersed outside of cell junctions **(Fig. 4G, Inx1& Inx3 rows, I, Inx2 row**). These data indicate that each cell type has a specific set of Innexins contributing to gap junction plaques.

In summary, our data highlight the unique roles of Innexins in regulating retinal wave activity through the formation of gap junction plaques. All retinal support cells require Inx2 for the formation of gap junction plaques that contain Inx1-3, whereas Inx3 is only required in interommatidial cells. Inx1 has a minor impact on its own but compensates for the absence of Inx3 in cone cells and primary pigment cells. When gap junction plaques were disrupted, the corresponding wave propagation pathway was also disrupted **(Fig. 4R**), illustrating the importance of precise Innexin combinations in plaques for directing waves across distinct retinal cell types.

### Retinal waves regulate Myosin II-driven apical remodeling that scales with ommatidial size

The ventro-anterior bias in retinal waves corresponds to a region with larger ommatidia **(Fig. 5A, E**), established by 24 hAPF and persisting through the wave period (**Fig. S5A-C**). These larger ommatidia showed stronger and prolonged calcium activity (**Fig. S5B, C**). To understand how this differential wave activity might influence the development of ommatidia in this region, we compared ommatidia before and after the waves **(Fig. 5F, G, M-P**). Prior to the waves, larger ommatidia displayed wider interommatidial gaps formed by the apical domains of interommatidial cells **(Fig. 5F top & bottom, M-P**). The 20% largest ventro-anterior ommatidia had wider boundaries of interommatidial cells than the 20% smallest dorso-posterior ommatidia **(Fig. 5O, P**). After the waves, interommatidial gaps contracted throughout the retina (**Fig. 5O, P**), whereas ommatidial size slightly increased (**Fig. S5F**). Larger ommatidia with initially wider gaps exhibited stronger boundary contraction (**Fig. 5O, P**). As a result, interommatidial boundaries became uniformly narrow across the retina (**Fig. 5G, P**), despite the regional variability in ommatidial size (**Fig. S5C**). Multi-color Flp-out-based single-cell labeling confirmed this contraction arose from apical morphogenesis of interommatidial cells (**Fig. S5G-P**).

**Fig. 5.**
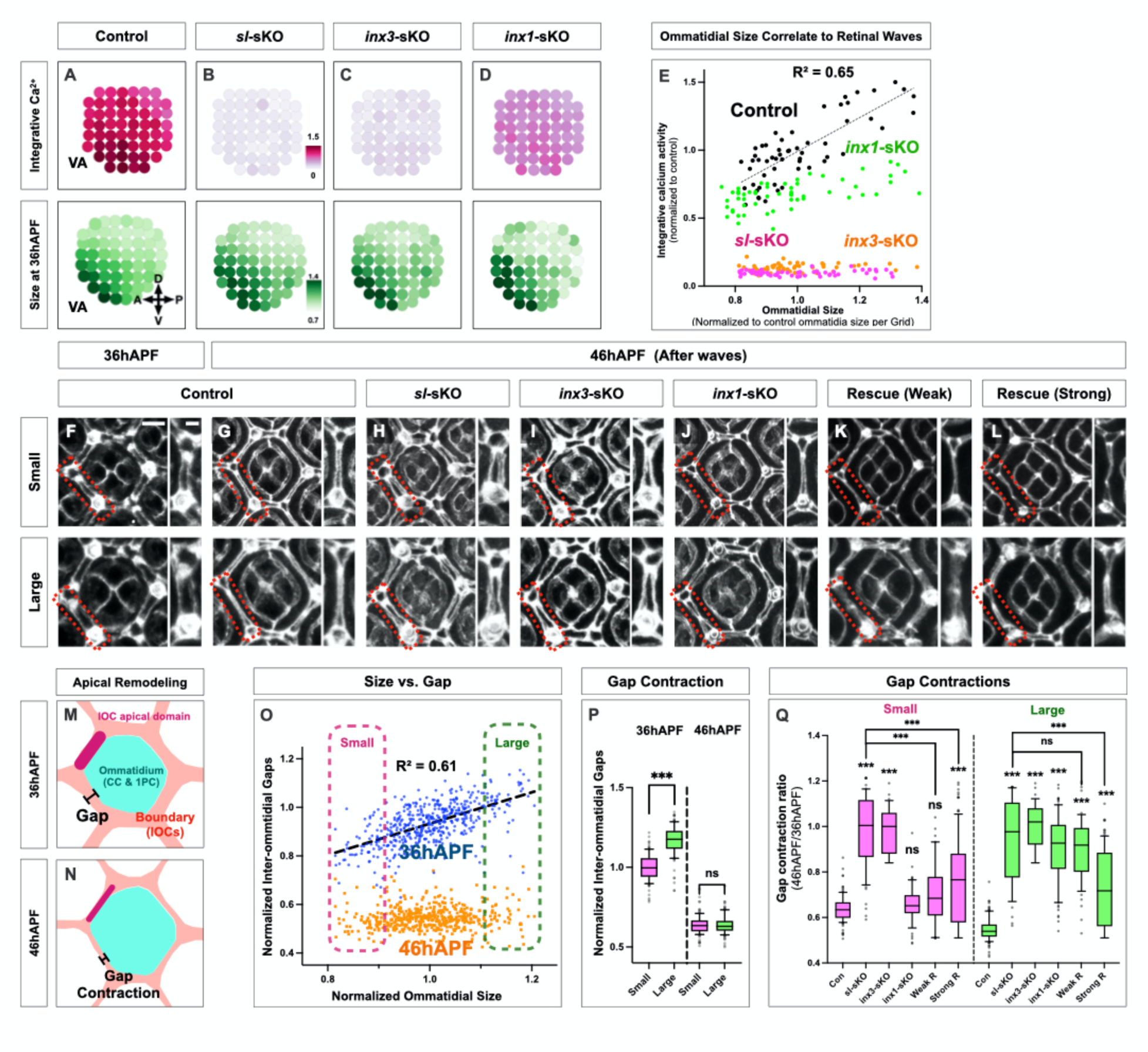
Retinal wave activity coordinates size-dependent ommatidial remodeling (A–Q). **5A–D.** Ommatidial 8×8 grids showing integrated calcium activity from retinal waves (**top**) and ommatidial size distributions (**bottom**) in control and mutants. Average intensity of integrative calcium activity and mean ommatidial size per grid point are color-coded in magenta and green, respectively. n ≥ 11 retinas. **5E.** Scatter plot of normalized ommatidial size (X) vs calcium signals (Y). Each dot represents a single grid point from (**A–D**). Genotypes are color-coded. The black line shows regression in control. **5F–L.** Dlg1 immunostaining of single ommatidia at 36 hAPF and 46 hAPF. **Tops** show small ommatidia (0–20%, dorsal-posterior region) and **bottoms** show large ommatidia (80–100%, ventral-anterior region). Red insets highlight interommatidial boundary gaps, enlarged at right. Scale bars: 5 μm **(F–L)**, 2 μm (insets). **5M, N.** Diagrams of ommatidial apical remodeling at 36 hAPF **(M)** and 46 hAPF **(N)**. Measurement bars indicate interommatidial gap size at boundaries. **5O.** Scatter plot of ommatidial size (X) vs gap size (Y) at 36 hAPF (blue) and 46 hAPF (red) from **(F–L)**. Each dot represents a single ommatidium. Values are normalized to the average of 36 hAPF. The regression line is shown for 36 hAPF. n ≥ 492 ommatidia from ≥ 16 retinas per stage. Dashed boxes indicate size groups for **(Q)**. **5P.** Quantification of interommatidial gaps in small and large size groups from **(O)**. n ≥ 101 ommatidia from ≥ 16 retinas per stage. ns: p ≥ 0.05, ***p < 0.001. Error bars: SD. **5Q.** Quantification of gap contraction ratio between 36 hAPF and 46 hAPF in small (magenta) and large (green) ommatidia from **(F, G)**. The Y-axis shows the ratio of gap size at 46 hAPF to the average at 36 hAPF for each size group and genotype. n ≥ 36 ommatidia from ≥ 16 retinas. ns: p ≥ 0.05, ***p < 0.001. Error bars: SD.

To assess whether retinal waves were required for this size-dependent contraction of ommatidial boundaries, we analyzed mutants with disrupted wave activity, either across all retinal cells (*sl* somatic knockout retinas) or specifically in interommatidial cells (*inx3* somatic knockout retinas) **(Fig. 5B, C, E**). Before retinal waves, ommatidial size distribution and boundary morphology were similar to control in both retinas **(Fig. 5B, C bottoms, H, I tops**). However, after retinal waves, boundary contraction failed in both mutants, resulting in ommatidial boundaries that remained immature and wider, independent of ommatidial size **(Fig. 5H, I bottoms, Q, Detailed in S6A**), suggesting that retinal waves are essential for ommatidial boundary contraction.

To further test how retinal waves coordinate boundary contraction in relation to ommatidial size, we analyzed *inx1* somatic knockout, which exhibit uniformly low calcium activity across all ommatidia, uncoupling calcium signaling from ommatidial size **(Fig. 5D, E**). As a result, boundary contraction failed selectively in the larger ventral-anterior ommatidia, while smaller ommatidia still showed proper contraction **(Fig. 5J, Q, S6A**). This suggests that wider initial boundaries require proportionally stronger wave activity to achieve uniform boundary maturation.

To directly test whether retinal waves drive apical boundary contraction proportionally to their activity, we manipulated calcium activity using the heat-activated calcium channel TrpA1 (*72, 73*). In *sl* somatic knockout, weak heat induction of TrpA1 (15 min) during the endogenous wave period (37 hAPF) only rescued boundary contraction in smaller ommatidia, while larger ommatidia failed to fully contract **(Fig. 5K, Q, S6A**). In contrast, stronger calcium induction (40 min) rescued boundary contraction in larger ommatidia **(Fig. 5L bottom, Q, S6A**) but caused abnormal contraction in smaller ommatidia, resulting in narrow and wrinkled boundaries **(Fig. 5L bottom**). These results suggest that the higher intensity of calcium signaling due to more frequent retinal waves drives stronger apical contraction and ensures uniform ommatidial organization.

IP_3_R-mediated calcium release activates myosin light-chain kinase, which phosphorylates non-muscle myosin II to form an actomyosin network, promoting contractility (*74, 75*). To examine Myosin II activation in response to calcium, we performed time-lapse imaging of retinas expressing GFP-tagged Sqh (Myosin II regulatory light chain). The Sqh-GFP signal was observed primarily as puncta of varying sizes on the apical side of the retina (**Fig. 6A, S6B, C**), indicating discrete actomyosin foci representing localized contractile activity (*76*). We quantified these dynamics by calculating the coefficient of variation of Sqh-GFP intensity along interommatidial boundaries (*77*). During retinal waves, this coefficient progressively increased, indicating the formation of larger and more organized actomyosin clusters within the interommatidial boundaries (**Fig. 6A-E, J**), consistent with wave-driven apical contraction of interommatidial cells. Notably, Sqh-GFP aggregation was stronger in interommatidial cells from larger ommatidia (**Fig. 6A-E bottoms, J**), supporting our model that higher local calcium signals drive greater contractile forces leading to robust apical contraction of interommatidial cells. Actomyosin network formation was markedly impaired in both *sl* and *inx3* somatic knockout retinas (**Fig. 6G, H, K**), while *inx1* somatic knockout retinas exhibited reduced Sqh-GFP clustering selectively in large ommatidia (**Fig. 6I bottom, K**).

**Fig. 6.**
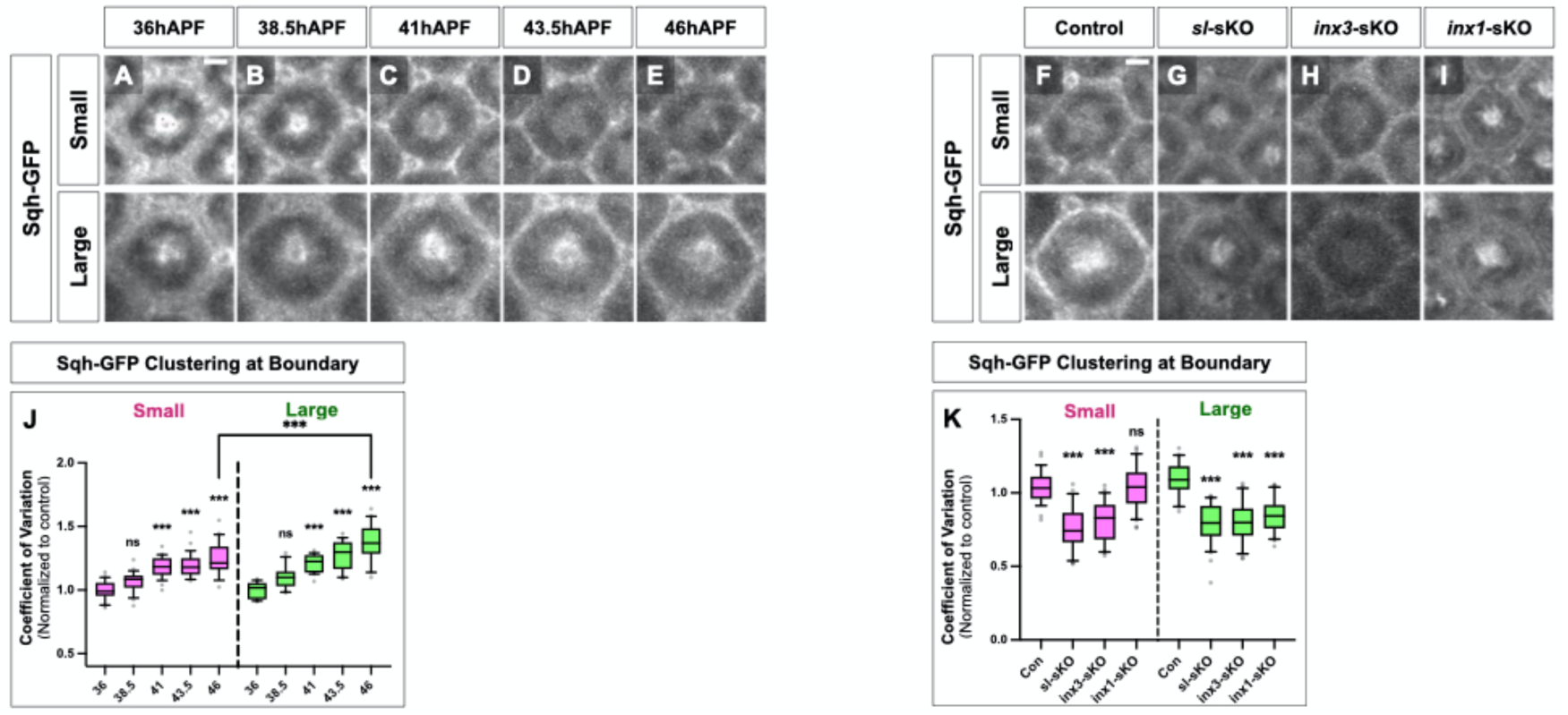
Retinal wave activity correlates with size-dependent Myosin II clustering (A–K). **6A–E.** Time-lapse images of single ommatidia from 36–46 hAPF at 2.5-h intervals showing Sqh-GFP puncta. **Tops** show small ommatidia, and **bottoms** show large ommatidia. Scale bar: 5 μm **(A–E)**. **6F–I.** Sqh-GFP puncta in single ommatidia at 46 hAPF. **Tops** show small ommatidia, and **bottoms** show large ommatidia. Genotypes indicated. Scale bar: 5 μm **(F–I)**. **6J.** Quantification of the coefficient of variation of Sqh-GFP within interommatidial boundaries from **(A–E)**. n ≥ 25 ommatidia from ≥ 5 retinas per genotype. ns: p ≥ 0.05, **p < 0.01, ***p < 0.001. Error bars: SD. **6K.** Quantification of the coefficient of variation of Sqh-GFP within interommatidial boundaries at 46 hAPF from **(F–I)**. n ≥ 22 ommatidia from ≥ 4 retinas. ns: p ≥ 0.05, ***p < 0.001. Error bars: SD.

To examine the temporal relationship between calcium activity and structural remodeling, we overlaid retinal wave timing, MyoII-GFP clustering, and interommatidial cell boundary contraction (**Fig. S6D**). Whereas minor morphological changes occurred during retinal waves, major structural remodeling was only complete by 46 hAPF, suggesting a temporal lag of ∼1–5 hours between the initial Ca²⁺ rise and detectable morphological changes.

Altogether, these results reveal that retinal waves coordinate apical contraction by scaling actomyosin network assembly proportionally to ommatidial size and boundary width, ensuring effective boundary contraction for uniform retinal morphogenesis.

### Retinal Wave Activity Ensures Eye Uniformity in Adults

To determine the functional role of retinal waves, we investigated how disrupted retinal waves during the mid-pupal stage affect adult retinal architecture. By 72 hAPF, the apical domains of interommatidial cells establish distinct ommatidial boundaries (**Fig. 7A, C** red arrow), acting as the basal footings of the corneal lens (**Fig. 7B**). In the adult eye, these footings are also critical for forming pseudocones, transparent extracellular structures positioned between the lens and photoreceptor neurons **(Fig. 7H, I**) (*78*), thereby ensuring precise light transmission to **PRs**.

**Fig. 7.**
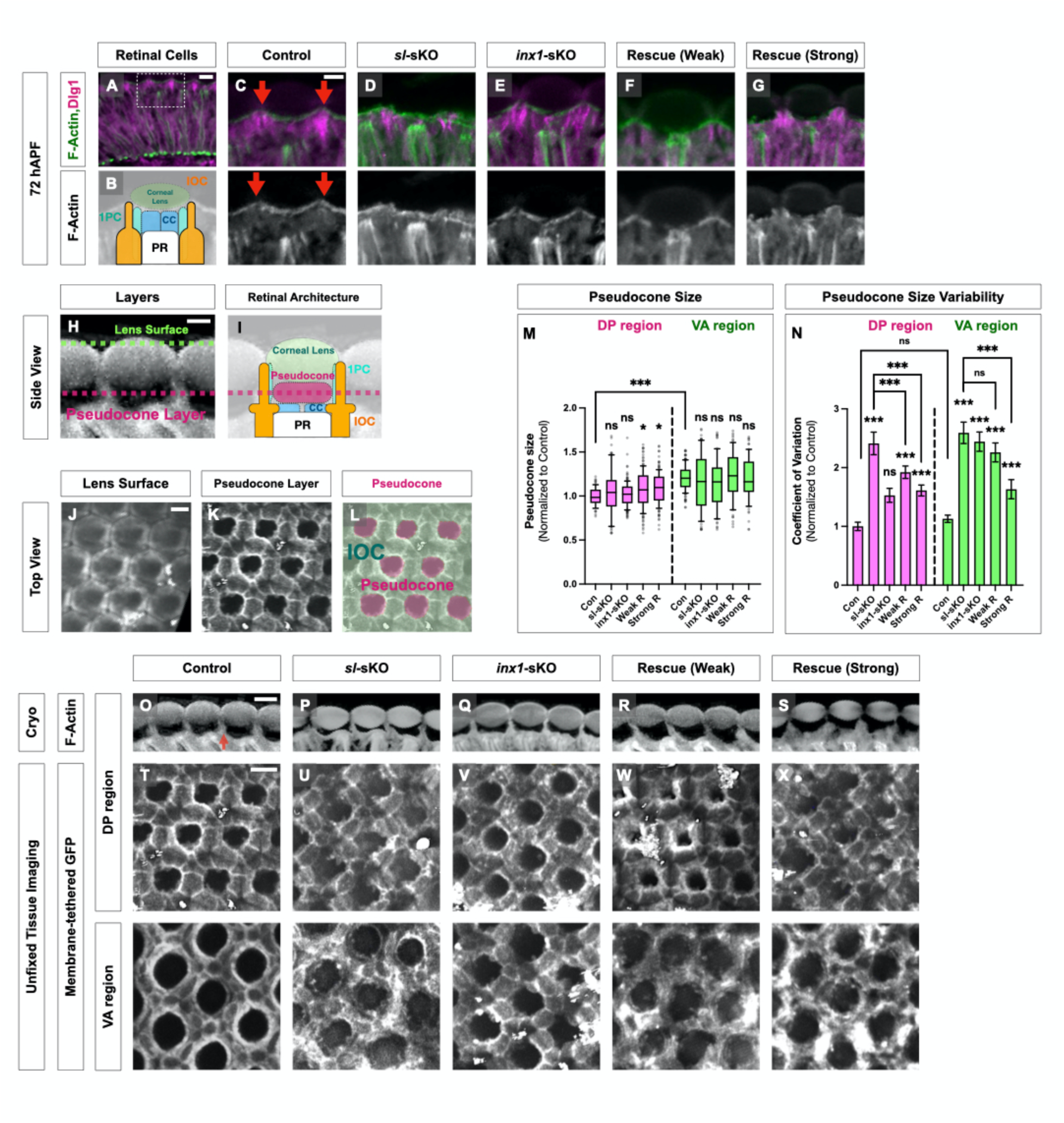
Retinal wave activity ensures adult eye uniformity (A–X). **7A.** Retinal surface at 72 hAPF in cryosection, labeled with phalloidin (green/gray) and Dlg1 (magenta). Red arrows in **(C)** show lens footings. **(A)** shows the whole section with an inset of apical surface **(C)**. **(B)** shows a diagram of retinal configuration. The dorsal-posterior region was selected. Z-stack: 5 μm. Scale bars: 10 μm **(A)**, 5 μm **(B–G)**. **7H, I.** Diagrams of adult retinal surface from side view, corresponding to **(O–S)**. Scale bar: 5 μm (H, I). **7J–L.** Adult retinal surface from top view. **(J, K)** show images, and **(L)** shows a diagram with pseudocones (magenta) and IOCs (green). Scale bar: 10 μm **(J–L)**. **7M.** Quantification of pseudocone size from **(T–X)** normalized to control. n ≥ 70 pseudocones from ≥ 15 retinas. ns: p ≥ 0.05, *p < 0.05. Error bars: SD. **7N.** Variability of pseudocone size from **(T–X)** measured by coefficient of variation. n ≥ 70 pseudocones from ≥ 15 retinas. Permutation test used. Error bars: SD of bootstrap resampling. ns: p ≥ 0.05, *p < 0.05, ***p < 0.001. **7O–S.** Adult retinal surface from side view. Cryosections were labeled with phalloidin. Red arrows in **(O)** show lens footings. Dorsal-posterior region selected. Z-stack: 5 μm. Scale bar: 10 μm **(O–S)**. **7T–X.** Adult retinal surface from top view. **Tops** show dorsal-posterior region, and **bottoms** show ventral-anterior region. Two-photon images of unfixed adult retinas were labeled with membrane-tethered GFP at the pseudocone layer **(L)**. Z-stack thickness: 10 μm. Scale bar: 10 μm **(T–X)**.

We quantified spatial organization of these structures in a single plane at the pseudocone layer (**Fig. 7H-L**). In wild type, pseudocone size correlated with ommatidial size: the ventro-anterior region (**VA**) contained consistently larger pseudocones than the dorso-posterior region (**DP**) **(Fig. 7T top and bottom, M**). In contrast, *sl* somatic knockout retinas showed severely disrupted pseudocone organization, although the outer lens surface appeared mostly normal **(Fig. S3F)**. Lens footings at 72 hAPF were defective as anticipated from earlier interommatidial cell defects (**Fig. 7D**), resulting in irregular pseudocone arrangements in adult retinas **(Fig. 7P, U**). Pseudocone sizes remained comparable (**Fig. 7M**), but their variability increased in the dorso-posterior and ventro-anterior regions (**Fig. 7U, N**). These results show retinal waves are essential for uniformly structured pseudocones by properly maturing interommatidial cells during mid-pupal development.

Mutants with varying retinal wave activity further supported this conclusion. In *inx1* somatic knockout retinas, lens footings and pseudocones remained normal in the dorso-posterior region, while variability greatly increased in the ventro-anterior region (**Fig. 7Q, V, N**). TrpA rescue experiments in *sl* somatic knockout retinas revealed regional differences: Weak rescue (15-minute heat induction) selectively improved lens footing structure and pseudocone uniformity in the dorso-posterior region but not in the ventro-anterior region (**Fig. 7R, W, N**). Strong rescue (40-minute heat induction) improved pseudocone uniformity in both dorso-posterior and ventro-anterior regions (**Fig. 7S, X, N**), although lens footings in the dorso-posterior region remained defective (**Fig. 7S, X**), matching interommatidial cell defects seen in smaller ommatidia at 46 hAPF (**Fig. 5L**).

Overall, these results demonstrate that retinal waves coordinate calcium-driven apical morphogenesis in proportion to ommatidial size, ensuring uniform retinal architecture essential for precise visual function. This highlights how synchronized calcium signaling directs tissue patterning (**Fig. S6E**).

## DISCUSSION

We described here the intricate dynamics of calcium activity due to retinal waves during early *Drosophila* retinal development and showed how it shapes retinal architecture by controlling the maturation of interommatidial cells. This provides insight into how directed synchronized calcium activity orchestrates structural patterning in tissues.

Regional variations in ommatidial size are evolutionarily conserved across fly species: Larger ventral ommatidia have larger lenses that enhance light capture from images of the ground and improve visual sensitivity, whereas smaller dorsal ommatidia increase resolution (*39, 40*). Despite this regional heterogeneity in ommatidial size, interommatidial boundaries remain uniformly spaced across the *Drosophila* retina and maintain a smooth and regular optical surface, ensuring proper lens alignment in the adult eye. Retinal waves mediate compensatory interommatidial cell maturation, ultimately normalizing interommatidial spacing across the retina.

In the *Drosophila* visual system, three phases of synchronized calcium activity shape development: two non-neuronal retinal waves are linked to tissue morphogenesis while neuronal synchronized activity is critical for synaptic wiring (*33, 62, 79, 80*). Importantly, early non-neuronal activity (early waves) precedes neuronal activity, highlighting a developmental sequence across different cell types. A comparable sequence is observed in vertebrates, where gap junction–mediated calcium waves in the retinal pigment epithelium of the chick eye and radial glia of the rodent neocortex precede synchronized neuronal activity of corresponding tissue during synaptogenesis (*13, 14, 21*). This progression from non-neuronal to neuronal calcium signaling positions *Drosophila* as a valuable model for studying fundamental principles of nervous system development.

The early retinal waves we observed occur at a precise stage, requiring coordination with other developmental events. This activity is initiated via the Cad96Ca RTK, which suggests that an extrinsic signal activating it determines the timing of retinal waves. While the Cad96Ca ligand is unknown, recent studies suggest that juvenile hormone can directly activate Cad96Ca in cotton bollworm and *Drosophila* S2 cells (*81*). Although the canonical juvenile hormone pathway involves the nuclear receptors Met and Gce that regulate juvenile hormone-responsive genes, Cad96Ca might be a membrane receptor for juvenile hormone, initiating rapid signaling independent of nuclear transcription. Systemic juvenile hormone amounts exhibit multiple peaks, including one at mid-pupal stages coinciding with the retinal wave period (*82*), raising the possibility that juvenile hormone directly activates Cad96Ca to trigger initial calcium activity of retinal waves.

Despite inherent differences in ommatidial size, proper visual function relies on a uniformly shaped pseudocone and lens in each facet. Optically, any mismatch between facet geometry and focal length broadens the point-spread function, reducing local contrast and spatial resolution (*83, 84*). Although we have not yet assessed visual performance directly, our structural observations imply that such mismatches could produce a mosaic of slightly blurred and sharper retinal areas. Future studies could therefore combine quantitative, high-resolution analyses of adult eye architecture with region-specific behavioral assays such as optomotor or luminance tests (*39, 85–87*). Such approaches may clarify the functional impact of irregular ommatidial packing and uncover how retinal calcium waves more broadly shape adult eye architecture and visual function.

Overall, our findings reveal three previously unappreciated features of calcium-based signaling during tissue morphogenesis. First, we identify Cad96Ca, an RTK, as a trigger of calcium activity. Second, we show that wave propagation is directed by a combinatorial “Innexin-code” rather than simple anatomical connectivity. Third, we demonstrate that calcium signaling scales with epithelial unit size to ensure proportional morphogenetic remodeling.

These insights suggest a general framework by which calcium-based intercellular communication orchestrates precise tissue patterning across diverse systems.

## Supporting information

All Movies

## ACKNOWLEDGMENTS

We thank the members of the Desplan lab for helpful discussions and manuscript feedback; J. Malin, S. Cordoba, and D. Chen for advice on manuscript preparation; N. Ghosh and J. Treisman for technical assistance with cryosectioning; M. Hoch and A. Borst for antibodies; P. Phelan for *shakB²* fly stocks; and M. Courgeon for plasmids. We used stocks obtained from the Bloomington Drosophila Stock Center (NIH P40OD018537). This work also benefited from antibodies and *Drosophila* lines generated with support from the CGSB at NYU Abu Dhabi.

## Funding

This work was supported by NIH grants EY13010 and EY017916 (C.D.), NIH fellowship F32EY027682 (B.J.C.), the MacCracken Fellowship at New York University (Y.-C.C.), a NYSTEM institutional training grant (C322560GG) (Y.-C.C.), and a scholarship to study abroad from the Ministry of Education, Taiwan (Y.-C.C.).

## Author contributions

B.J.C. and C.D. conceived the project, analyzed the data, and wrote the manuscript. Y.-C.C. performed quantitative analysis of calcium imaging data. B.J.C. performed all experiments.

## Competing interests

The authors declare no competing interests.

## Data and materials availability

All datasets generated during this study are deposited in the Dryad repository (*90*). Antibodies and fly lines will be distributed upon request. All analysis code is available at Zenodo (*91*).

## Material and Methods

### A complete list of genotypes and reagents is provided in Supplementary Materials and Tables

#### Drosophila melanogaster Strains and Husbandry

All *Drosophila melanogaster* stocks used are listed in the **<u>Materials</u>**. Flies were reared on standard cornmeal medium under 12-hour light/dark cycles at 25°C, unless specified otherwise. Only male flies were analyzed to take advantage of enhanced efficacy of somatic knockout of major target genes (*sl*, *inx1*, *inx2*) located on the X chromosome. Pupal staging was performed by marking white pupae adhered to the dry wall of the vial, defined as 0 hAPF (hours after puparium formation). Pupae exhibiting significantly smaller body size were excluded from all experiments. Detailed genotypes corresponding to each figure are provided in **Table S2**.

#### Generation of Transgenic Animals

Transgenic constructs including longGMR-Cas9, customized guide RNAs, and GMR-GCaMP6s were generated by GenScript (Piscataway, NJ, USA). Insertions were synthesized into appropriate plasmid backbones (**see Table S3** for plasmid details). All transgenes were injected into corresponding attP sites by BestGene (Chino Hills, CA, USA) (**see <u>Materials</u>**for transgene information). For guide RNAs, we used the pCFD5_w vector (*57*) with the U6:3 promoter to ubiquitously express six tandem guide RNAs, consisting of two duplicated sets targeting three selected sites within the first and second common exons (see **Table S3** for guide RNA details). This configuration produced a strong somatic knock-out effect across all tested targets, exemplified in **Fig. S4U.**

#### Live Imaging of Intact Pupal Retina

Pupae within the selected developmental window were gently attached to double-sided tape (Scotch) on a slide in the desired position. The pupal case surrounding the head was minimally removed using fine-tip forceps without touching or pulling the pupal body. Samples were imaged using dry-objective lenses with a semi-open chamber surrounded by wet Kimwipes, utilizing either a Leica SP8 confocal microscope or an Olympus FV1000MPE2 two-photon microscope. Imaging was performed using 488 nm (confocal, Leica SP8) or 920 nm (two-photon, Olympus FV1000MPE2; Mai Tai Ti:Sapphire Lasers) excitation lasers.

All images were acquired with a **10x Leica dry-objective (confocal)** or a **20x Olympus dry-objective (two-photon).** For confocal imaging, the entire retina was scanned using a **10x objective with 3∼4x zoom.** For two-photon imaging, the flattest local area of each mounted retina, between the central and dorsal retinal regions, was scanned using a **20x objective with 8x zoom.** Images were collected using **LAS-X (Leica)** at **1024 × 800-pixel resolution** or **Fluoview (Olympus)** at **512 × 512-pixel resolution**. Laser power was adjusted to the minimum necessary to avoid potential phototoxicity and photobleaching.

Images displaying ectopic local or global increases in basal calcium signals during scanning (indicating potential phototoxicity) or significant reductions in basal calcium signals (indicating abnormal photobleaching) were excluded from further analysis.

Each scanning session lasted up to **12 hours for any analysis** to avoid potential phototoxicity. After successful imaging, animals were returned to standard fly food with wet Kimwipes.

Successfully scanned animals that were returned to food developed normally into adults without gross morphological defects.

#### Pan-retinal expression of genetically encoded calcium indicator and Cas9

Calcium imaging was performed using longGMR-GAL4 to drive UAS-GCaMP6s unless otherwise noted. While longGMR-GAL4 is primarily photoreceptor-specific at larval stages, it broadly expresses in all retinal cells, including non-neuronal support cells, by 24 hAPF. Therefore, GMR-GAL4 was used for early developmental stages (12–24 hAPF; **Fig. 1C, S1A**), and longGMR-GAL4 was used as a pan-retinal driver for later stages throughout this study.

Similarly, longGMR-Cas9 does not express in non-neuronal support cells at larval stages, allowing bypass of early developmental requirements for target genes such as *inx1*, *inx2*, *inx3*, and *sl*.

#### Unfixed Tissue Imaging of Adult Retina

Adult animals younger than 5 days old were sacrificed and mounted at the center of a petri dish (10 × 35 mm, Thermo Scientific) using nail-polish (Amazon) with cold distilled water. Samples were positioned such that the targeted eye region (dorsal-posterior or ventral-anterior regions) was placed on the flat surface. Imaging was performed using an Olympus two-photon microscope with 920 nm excitation and an Olympus 25x water-immersion objective. Images compromised by air-bubble entrapment, preventing clear visualization, were excluded from analysis.

#### Confocal Imaging of Fixed Tissue

Fixed samples were imaged using a Leica SP8 confocal microscope or a Leica Stellaris confocal microscope with STED module (**Fig. 4A-D**) with a 63x or 100x (STED) glycerol-immersion objective lens. Samples were positioned such that the targeted eye region (dorsal-posterior or ventral-anterior regions) was placed on the flat surface. Images were analyzed using FIJI (ImageJ) and Imaris for further analysis.

#### Cryosection

Adult or pupal heads were fixed, processed through a sucrose gradient, embedded in OCT, cryosectioned at 12 μm, and postfixed. We focused exclusively on the dorsal-posterior region, as its proximity to the flat center of the adult eye allows more robust sectioning (**Fig. 7A-G, O-S**). Detailed protocols are provided in supplementary references (*88*).

#### Immunohistochemistry

Fly pupal retina and optic lobes were dissected, fixed, blocked, and stained using standard immunohistochemistry protocols. Full details are provided in supplementary references (*89*).

The following primary antibodies were used for immunofluorescence: mouse anti-Dlg1 (1:200, 4F3, DSHB), rabbit anti-HA (1:200, Cell Signaling Technologies), rat anti-FLAG (1:200, Novus), mouse Anti-V5-tag:DyLight550 (1:200, BioRad), guinea pig anti-inx1 (1:1k, GenScript), rabbit anti-inx2 (1:2k, M. Hoch), guinea pig anti-inx3 (1:1k, GenScript), rabbit anti-inx4 (1:1k, A. Borst), rabbit anti-inx5 (1:1k, A. Borst), rabbit anti-inx6 (1:1k, A. Borst), rabbit anti-inx7 (1:1k, A. Borst), rabbit anti-inx8 (1:1k, A. Borst), chicken anti-GFP (1:1k, EMD), rat anti-Ecad (1:10, DSHB, DCAD2), AlexaFluor660 conjugated Phalloidin (Invitrogen, 1:200), and AlexaFluor405 conjugated Goat Anti-HRP (Jackson ImmunoResearch, 1:100). Secondary antibodies were from Invitrogen and used at 1:200.

#### TrpA Heat Activation

Pupae at 37 hAPF were selected, attached to double-sided tape on glass slides, and the pupal case around the retinas was removed. Animals were then incubated in a dark 29°C incubator for the designated time, followed by transfer to a 25°C incubator with wet Kimwipes to maintain humidity. Samples were dissected at 46 hAPF (**Fig. 5**), 72 hAPF (**Fig. 7F, G**), and the adult stage (**Fig. 7R, S, W, X**) for further analysis. Animals showing obvious signs of cellular toxicity from heat induction, such as darker pigmentation or scarring, were excluded from experiments.

## QUANTIFICATION AND STATISTICAL ANALYSIS

### Quantification of Retinal Wave-Induced Calcium Activity

Calcium imaging data from SP8 confocal microscopy of whole retinas was quantified using a hybrid model integrating event detection and intensity measurement for improved sensitivity and accuracy. Time-lapse images were pre-processed in Fiji to obtain corrected F/F0 values. Photobleaching was corrected by exponential decay fitting in Fiji under consistent exposure settings. ROIs for individual ommatidia were defined in a semi-automatic manner for further analysis in R. A two-component Gaussian mixture model (GMM) was applied to classify each ROI in each frame as “On” or “Off”, with thresholds semi-manually optimized to enhance detection accuracy. Total calcium activity was calculated by summing ΔF/F0 values exclusively during “On” states, reducing background noise and ensuring meaningful activity quantification. This approach improves sensitivity and accuracy by combining event frequency and intensity.

Calcium imaging data from two-photon microscopy with limited ROIs at single-cell resolution was quantified by integrating all ΔF/F0 intensity values to capture total calcium signal magnitude. Time-lapse images were pre-processed in Fiji to obtain corrected F/F0 values. All ΔF/F0 values across corresponding frames were summed to quantify total calcium activity. ROIs for individual cell types were defined semi-automatically. Retinal pathways were manually defined, considering sequential activation of calcium activity from adjacent cells.

### Imaris-based Morphological Visualization and Analysis

For morphological analysis (**Fig. 5-7**), confocal Z-stacks were imported into Imaris (Bitplane) and visualized in 3D view. Planes and viewing angles were selected based on the region of interest. Single-plane images with optical thickness of 5 μm (**Fig. 5**) or 10 μm (**Fig. 7**) were captured from 3D-rendered volumes for representative figures and morphological analysis.

Morphological quantification was performed on captured planes, as the structures are much larger than pixel scale and only minimally influenced by standard Imaris 3D rendering steps. Selection of optical sections was based on morphological criteria described in each quantification protocol below. For intensity quantification (**Fig. 6**), Imaris 3D view was used for ROI selection, but actual quantification was performed using raw voxel data. All images (**Fig. 5-7**) used for figure panels were resized post-capture to match the same physical scale across conditions. Scale bars were adjusted accordingly to reflect accurate physical dimensions.

### Quantification of Ommatidial Morphology (Fig. 5, 7)

For whole-retina ommatidial size analysis, retinas expressing longGMR-GAL4-driven membrane-tethered GFP were imaged in intact animals using a two-photon microscope at 0.3 μm voxel resolution. Full Z-stacks were used to reconstruct a 3D retinal surface, and individual ommatidial hexagonal areas were calculated using Imaris software, which semi-automatically detected the apical side of each hexagonal corner point.

For high-resolution analysis of ommatidial boundaries, dissected retinas immunostained with Dlg1 were flat-mounted to position the region of interest. Confocal Z-stacks were acquired using an SP8 microscope and imported into Imaris for 3D visualization. To quantify each ommatidium from the correct position, morphologically defined optical sections (5 μm thickness) were selected to capture the most apical tip of Dlg1 staining and were extracted from 3D-rendered views for analysis and figure panels (**Fig. 5F–Q**). In each ommatidium, the gap distance was measured between apical Dlg1 staining along a line connecting ommatidial centers.

Interommatidial gaps along the dorsal-ventral axis adjacent to 1PC/1PC junctions were excluded due to their consistently reduced apical domains compared to other orientations, and the average of the remaining four gaps was assigned per ommatidium.

*Innexin Intensity (Fig. S4W–Y)*

Fixed pupal retinas were stained with innexin antibodies and Dlg1. Images were acquired using an SP8 confocal microscope, and innexin intensity within interommatidial junctional Dlg1 was quantified.

### Coefficient of Variation of Sqh-GFP (*Fig. 6*)

Retinas expressing Sqh-GFP were imaged in intact animals using a two-photon microscope with a 0.4 μm voxel size to reconstruct the 3D retinal surface. Time-lapse series (**Fig. 6J**) were corrected for photobleaching using the exponential decay method in Fiji under consistent exposure settings. Images were analyzed using Imaris, with manual refinement to define ROIs corresponding to the apical interommatidial boundary areas. To quantify local clustering of Sqh-GFP signal, the coefficient of variation (**CV**) was calculated within these defined ROIs. All pixel intensity values within each ROI, based on raw voxel data, were extracted, and the CV was calculated as the ratio of the standard deviation to the mean intensity. Higher CV values indicate greater intensity heterogeneity, reflecting localized clustering or accumulation of fluorescence signal.

### Pseudocone Quantification (*Fig. 7M, N*)

Adult eye pseudocones were imaged using a two-photon microscope at pseudocone layers within targeted eye regions (dorsal-posterior or ventral-anterior). Using Imaris, an optical section was selected to capture a consistent pseudocone layer (**Fig. 7K**). Pseudocone areas were semi-automatically defined as empty spaces enclosed by interommatidial boundaries. For morphological quantification of pseudocones, image display settings were adjusted individually to improve boundary visibility. Measurements were based on shape and size, not signal intensity. Any pseudocones obscured by bubbles or debris were excluded from the quantification.

### Statistical Analysis

Male pupae at the same developmental stage within a 1-hour time window were randomly selected from fly vials for all experiments. Blinded analysis across different genotypes was not performed, as genotypes were distinguishable by the experimenter. All experiments were conducted with at least three biological replicates (retinas from three different animals) per genotype. Statistical comparisons between genotypes were performed using the Mann–Whitney U test for two-group comparisons and the Kruskal–Wallis test followed by Dunn’s multiple comparisons test for comparisons involving more than two groups. Linear regression analyses were performed using GraphPad Prism. Data are presented as mean ± standard deviation (SD) or standard error of the mean (SEM), as indicated in figure legends. To test statistical significance of CV differences between groups (**Fig. 7N**), we used a permutation test, which does not assume normality and is well suited for comparing variability metrics such as CV.

## Use of AI technologies

During manuscript preparation, the authors used ChatGPT Plus (OpenAI, GPT-4 and GPT-5, 2024–2025 versions) for language editing only. All authors reviewed and edited the text after use and take full responsibility for the content of the publication.

## Supplementary Materials

Figs. S1 to S6

Tables S1 to S3

Movies S1 to S3

## MOVIE TITLES AND LEGENDS

**Movie 1. Calcium dynamics of developing pupal retina from 12 hAPF to eclosion (A–I),** related to **Fig. S1A-K.** Representative calcium imaging of intact pupal retinas from 12–24 hAPF **(A),** 24–36 hAPF **(B),** 36–48 hAPF **(C),** 48–59 hAPF **(D),** 60–72 hAPF **(E),** 72–84 hAPF **(F),** 84 hAPF to eclosion **(G),** early retinal waves **(H),** and late retinal waves **(I)** using *UAS-GCaMP6s* driven by *GMR-GAL4* **(A)** or *longGMR-GAL4* **(B–I).** Imaging performed with confocal microscopy (488nm excitation), focusing on retina (**A-G**) or entire pupa (**H, I**). Original GMR-GAL4 is expressed in all retinal cells post-morphogenetic furrow, while longGMR-GAL4 is photoreceptor-specific during the L3 larval eye disc but becomes pan-retinal by 24 hAPF. Time-lapse imaging was acquired as Z-stacks at 2 min intervals. Imaging duration was limited to a maximum of 12 hours per pupa to minimize phototoxicity for any analysis.

**Movie 2. Retinal calcium waves of a single retina (A–D),** related to **Fig. 1D-G**. Representative retinal calcium wave recordings from a single retina of an intact pupa during early stage 1 **(A),** late stage 1 **(B),** stage 2 **(C),** and stage 3 **(D)** using *UAS-GCaMP6s* driven by *longGMR-GAL4*. Stages are defined in **Fig. 1H**. Imaging performed with confocal microscopy (488nm excitation). Time-lapse imaging was acquired as Z-stacks at 3 s intervals.

**Movie 3. Retinal calcium waves at single-cell resolution (A–D),** related to **Fig. 2I-L**. Representative recordings of retinal calcium waves from a single retina during early stage 1 **(A),** late stage 1 **(B),** stage 2 **(C),** and stage 3 **(D)** using *UAS-GCaMP6s* driven by *longGMR-GAL4.* Imaging performed with two-photon microscopy (920nm excitation). Time-lapse imaging was acquired as Z-stacks at 5 s intervals.

**Movie 4. Vertical pathways of retinal calcium waves (A-D),** related to **Fig. 2M-P, S2A, C–E.** Representative recordings of retinal calcium waves during stage 1 showing initial cone cell firing **(A),** cone cell to cone cell propagation **(B),** cone cell to cone cell to primary pigment cell propagation **(C),** and cone cell to cone cell to primary pigment cell to interommatidial cells propagation **(D).** Imaging performed with two-photon microscopy (920nm excitation). Z-stacks were acquired every 4 s. Apical and basal layers are color-coded as green (apical) and magenta (basal).

**Movie 5. Lateral pathways of retinal calcium waves**, related to **Fig. 2R, S, S2F**. Representative recordings of retinal calcium wave propagation across multiple ommatidia during stage 2 using *UAS-GCaMP6s* driven by *longGMR-GAL4.* Imaging performed with two-photon microscopy (920nm excitation). Z-stacks were acquired every 4 s. Apical and basal layers are color-coded as green (apical) and magenta (basal).

**Movie 6. Dual-color calcium imaging of ER-released calcium activity during retinal waves (A-C)**, related to **Fig. 3A-C**. Representative recording of retinal calcium waves during stage 2 using *UAS-ER-GCaMP6* and *UAS-jRGECO1a* driven by *longGMR-GAL4*. Imaging performed with confocal microscopy (488 nm and 561 nm excitation). Time-lapse imaging was acquired as Z-stacks at 1 sec intervals. Dual-color imaging with ER-targeted GCaMP (green) and cytosolic jRGECO1a (red) (**A**), ER-targeted GCaMP alone (**B**), and cytosolic jRGECO1a alone (**C**).

**Movie 7. Calcium imaging of IP_3_R clonal retinas during retinal waves, related to Fig. 3F**. Representative recording of retinal calcium waves using *UAS-GCaMP8s* driven *by longGMR-GAL4* in *IP_3_R* clonal retinas during stage 2. Imaging performed with two-photon microscopy (920nm excitation). Time-lapse imaging was acquired as Z-stacks at 5 s intervals.

**Movie 8. Calcium imaging of mutants affecting wave initiation (A–E),** related to **Fig. 3I-M**. Representative recordings of retinal calcium waves using *UAS-GCaMP6s* driven by *longGMR-GAL4* in control **(A),** *norpA* mutants **(B),** *sl* mutants **(C),** *sl*-sKO **(D),** and *cad96Ca*-sKO **(E)** during a 5-minute peak intensity window at stage 2. Imaging performed with two-photon microscopy (920nm excitation). Time-lapse imaging was acquired as Z-stacks at 5 s intervals.

**Movie 9. Calcium imaging of *inx* mutants during retinal waves (A–E),** related to **Fig. 4J-N**. Representative recordings of retinal calcium waves using *UAS-GCaMP6s* driven by *longGMR-GAL4* in control **(A),** *inx1*-sKO **(B),** *inx2*-sKO **(C),** *inx3*-sKO **(D),** and *inx1+3*-DsKO **(E)** during a 5-minute peak intensity window at stage 2. Imaging performed with two-photon microscopy (920nm excitation). Time-lapse imaging was acquired as Z-stacks at 5 s intervals.

## Materials

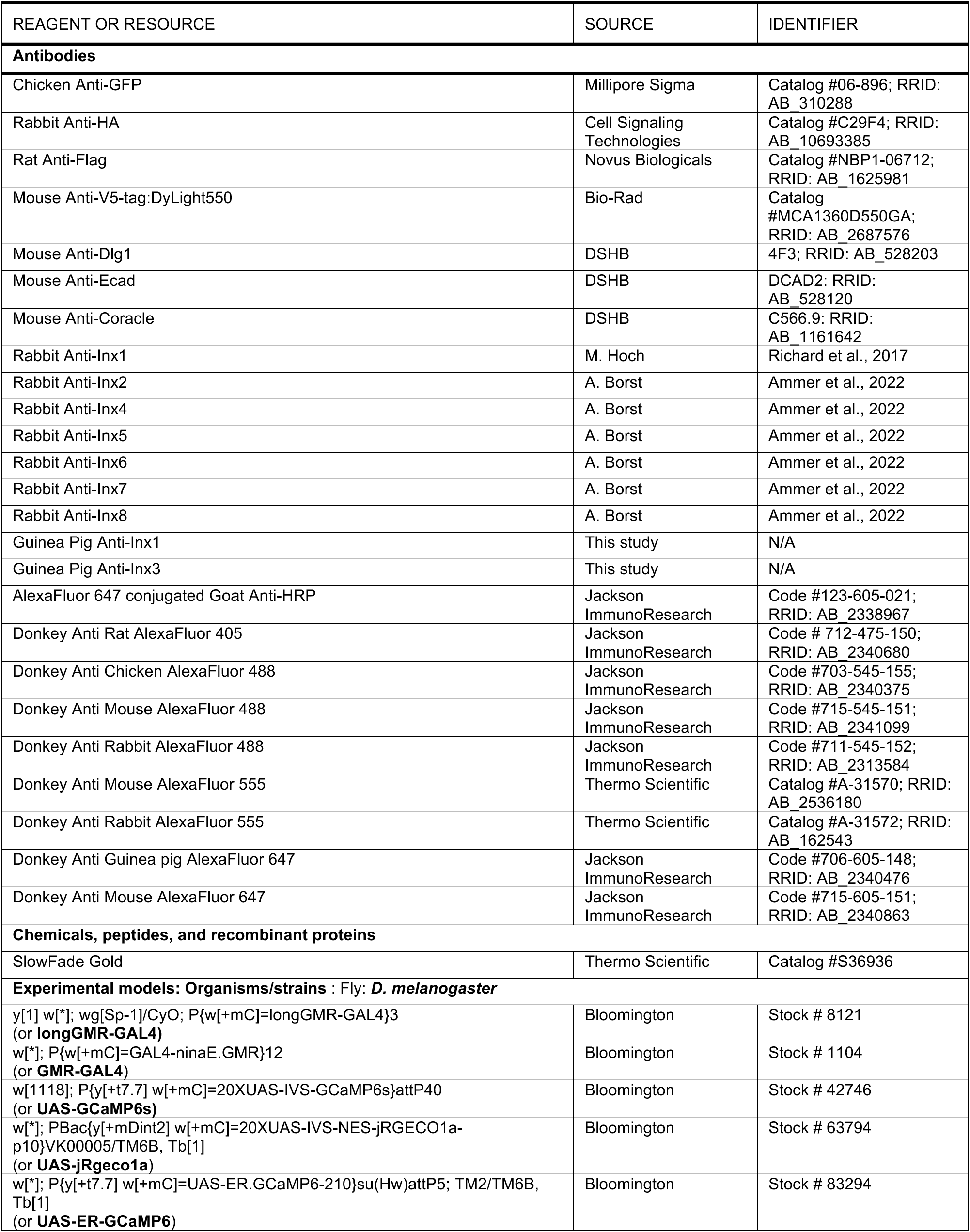
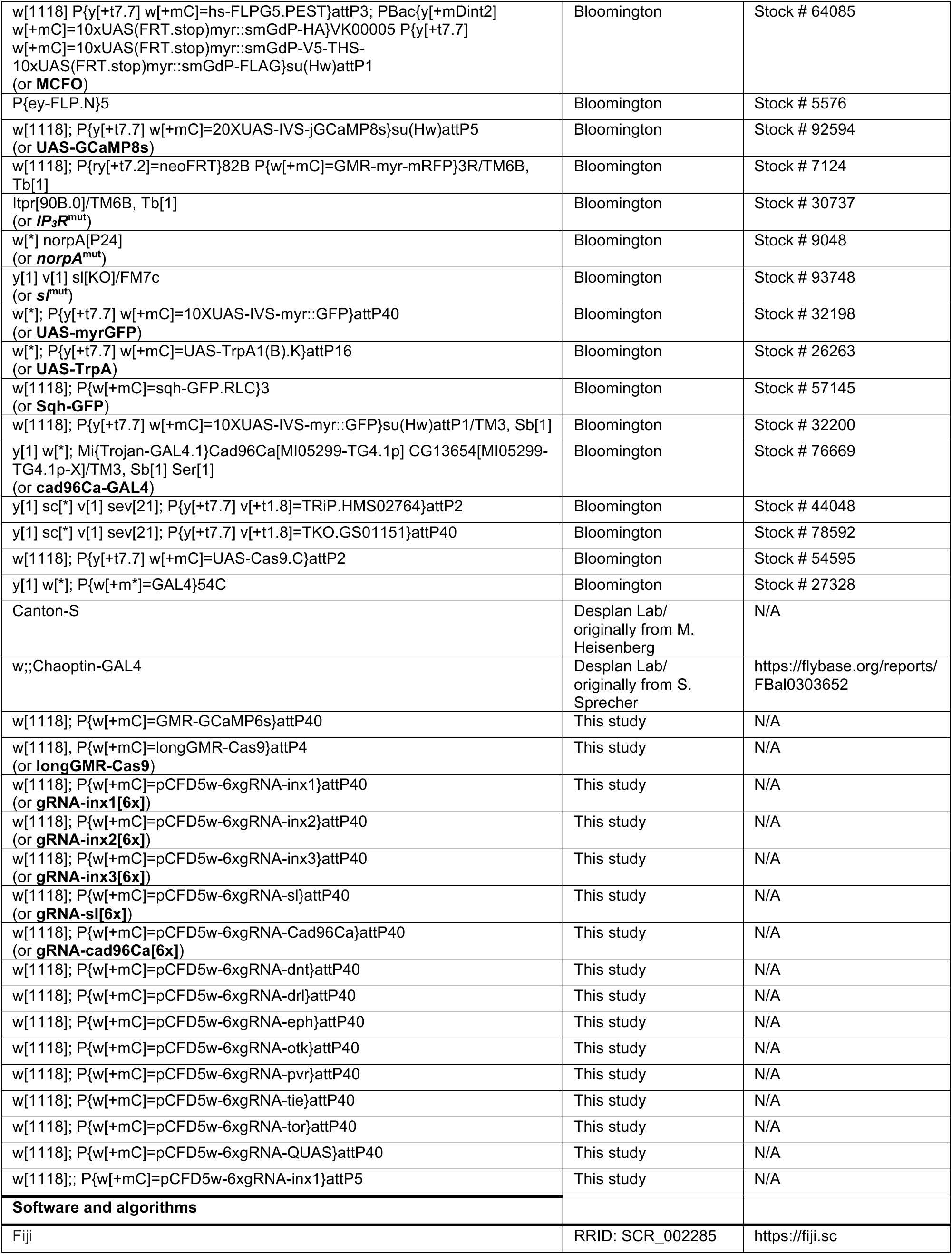
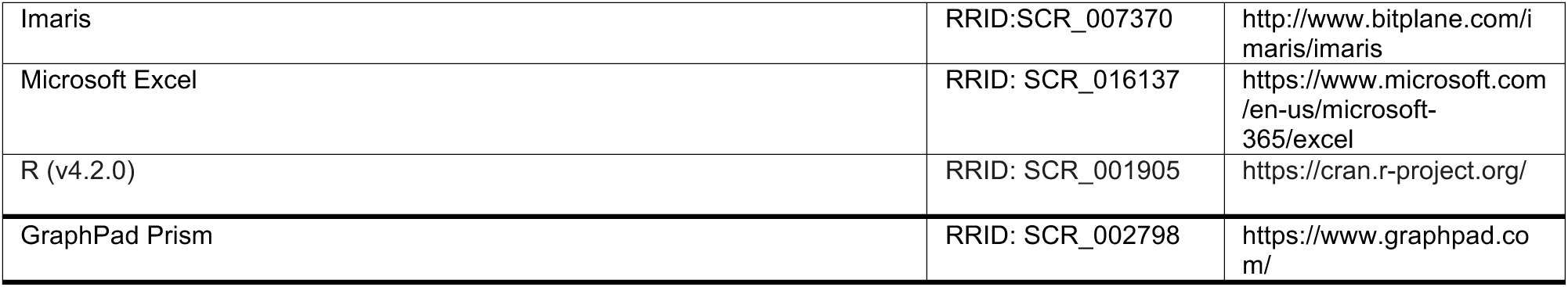

## Supplementary Table Legends

**Table S1. RTK screen for regulators of retinal waves, related to Fig. 3**.

Receptor tyrosine kinases expressed in the developing retina were tested by the indicated genetic manipulations. Screening was performed under the same conditions as (**Fig. 1C**) in a longGMR-GAL4>UAS-GCaMP6s background. At least five animals per condition were imaged continuously for ∼12 h between 36–48 hAPF, and gross retinal morphology was subsequently examined by immunostaining.

**Table S2. Experimental genotypes corresponding to figures.**

Genotypes used for each figure are listed with both the common label and the full genotype description.

**Table S3. Transgenic constructs and plasmid resources.**

Sequence information for newly generated transgenic constructs is provided, including longGMR-Cas9, customized guide RNAs, and GMR-GCaMP6s.

## Supplementary Movie Legends

**Movie S1. Bilateral retinal calcium waves**, related to **Fig. S1L.** Representative recording of bilateral retinal calcium waves from both eyes of a single animal during retinal waves using *UAS-GCaMP6s* driven by *longGMR-GAL4.* Imaging performed with confocal microscopy (488nm excitation). Time-lapse imaging was acquired as Z-stacks at 2-minute intervals.

**Movie S2. Calcium imaging of photoreceptor neurons during retinal waves**, related to **Fig. S3A, B.** Representative recording of retinal calcium waves using *UAS-GCaMP6s* driven by *Chaoptin-GAL4* during stage 2. Imaging performed with two-photon microscopy (920nm excitation). Z-stacks were acquired every 4 sec.

**Movie S3. Calcium imaging of *inx8* mutants during retinal waves**, related to **Fig. S4O, P.** Representative recording of retinal calcium waves using *UAS-GCaMP6s* driven by *longGMR-GAL4 in inx8* mutant retinas during a 5-minute peak intensity window at stage 2. Imaging performed with two-photon microscopy (920nm excitation). Time-lapse imaging was acquired as Z-stacks at 5 sec intervals.

---

**Fig. S1 (A–G) related to Fig. 1.**
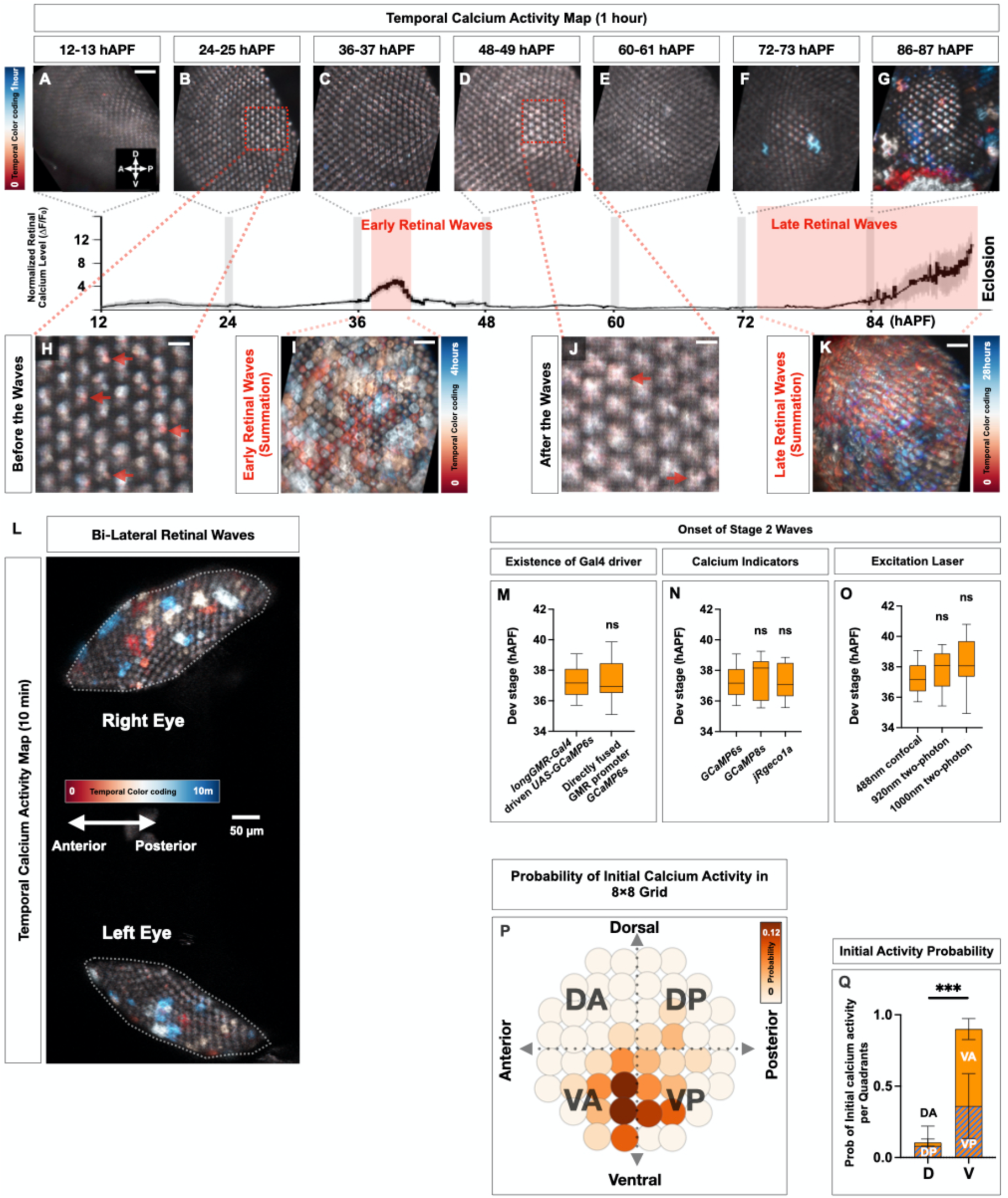
**S1A-K. Trace of average retinal calcium dynamics throughout pupal retinal development** (same as **1C**) with a representative color-coded temporal calcium activity map (1 hour) for each developmental time window (marked in gray window in the trace). **A-G** show temporal calcium activity maps at every 12-hour interval, with insets of **B, D** shown in **H, J** at higher magnification. **I, K** display the temporal activity maps for the entire early and late retinal wave periods, respectively. Calcium imaging was performed using *longGMR-GAL4* (or *GMR-GAL4* for 12-24hAPF) driving *UAS-GCaMP6s* and images taken at 2-minute intervals, with n ≥ 5 retinas analyzed per 12-hour time window. Scale bar: 10 μm **(A-G, I, K),** 2.5 μm **(H, J).** See also **Movie 1**. **S1L. Representative temporal calcium activity map across 10 minutes**, covering both eyes during stage 2 retinal waves, demonstrating bilateral wave synchronization. See also **Movie S1**. **S1M-O. Quantification of the onset of stage 2 retinal waves under different experimental conditions.** n ≥ 8 retinas. Calcium imaging was performed using indicated genotypes (**M**), *longGMR-GAL4* driving UAS-transgenes of indicated calcium sensors **(N),** or *UAS-GCaMP6s* **(O)**. ns: p ≥ 0.05, error bars: SD. **S1P**. **Ommatidial configuration grid showing spatial pattern of initial activity.** Each circle represents a group of ommatidia within an 8×8 grid along the dorsoventral (**DV**) and anteroposterior (**AP**) axes. Color indicates the probability of initial calcium activity (five first-triggering ommatidia per retina). Data pooled from 50 events across 10 control retinas. **S1Q**. **Quantification of initial calcium activity probability by quadrant** (from **Fig. S1P**) n = 50 events from 10 control retinas. ns: p ≥ 0.05; ***p < 0.001, error bars: SD.

**Fig. S2 (A–F) related to Fig. 2.**
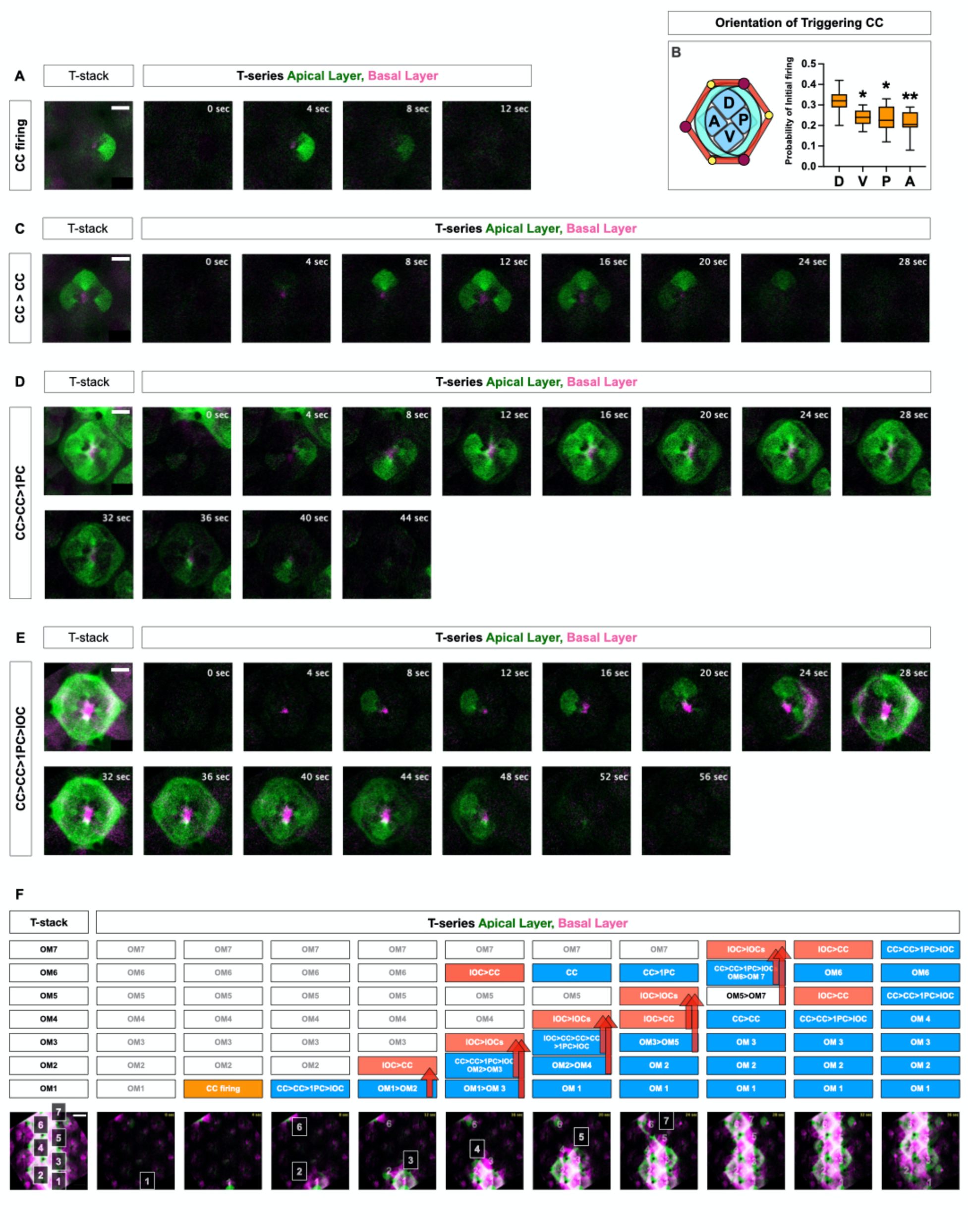
**S2A, C-E.** Representative calcium imaging of the vertical pathway of retinal waves, including single CC firing (**A**), CC to CC (**C**), CC to CC to 1PC (**D**), and CC to CC to 1PC to IOC (**E**). Integrative T-stacks and individual images of corresponding T-series (4-sec interval) are shown with color-coded layer-specific calcium signal as apical and basal layers in green and magenta, respectively. The apical layer signals mostly originate from CCs and 1PCs, while the basal layer signals mostly originate from IOCs. CC = cone cell, 1PC = primary pigment cell, IOC = inter-ommatidial cells. Scale bar: 5 μm **(A, C-E).** See also **Movie 4**. **S2B.** Quantification of initial calcium activity probability by cone-cell orientation n = 239 ommatidia from 10 control retinas. ns: p ≥ 0.05; *p < 0.05; **p < 0.01, error bars: SD. **S2F.** Representative retinal wave propagating across multiple ommatidia using both vertical and lateral pathways. Each box marks marked ommatidia (**OM1–7**) along the wave path where each wave-induced event is color-coded. Yellow indicates spontaneously emerging CC firing, blue indicates vertical propagation within those ommatidia, and red indicates target ommatidia activated by lateral pathways. The red arrow highlights a lateral pathway between two ommatidia. In the frame of 16 sec, OM6 is activated by other waves from outside the imaging area via the lateral pathway. Scale bar: 10 μm. See also **Movie 5**.

**Fig. S3 (A–L) related to Fig. 2 and 3.**
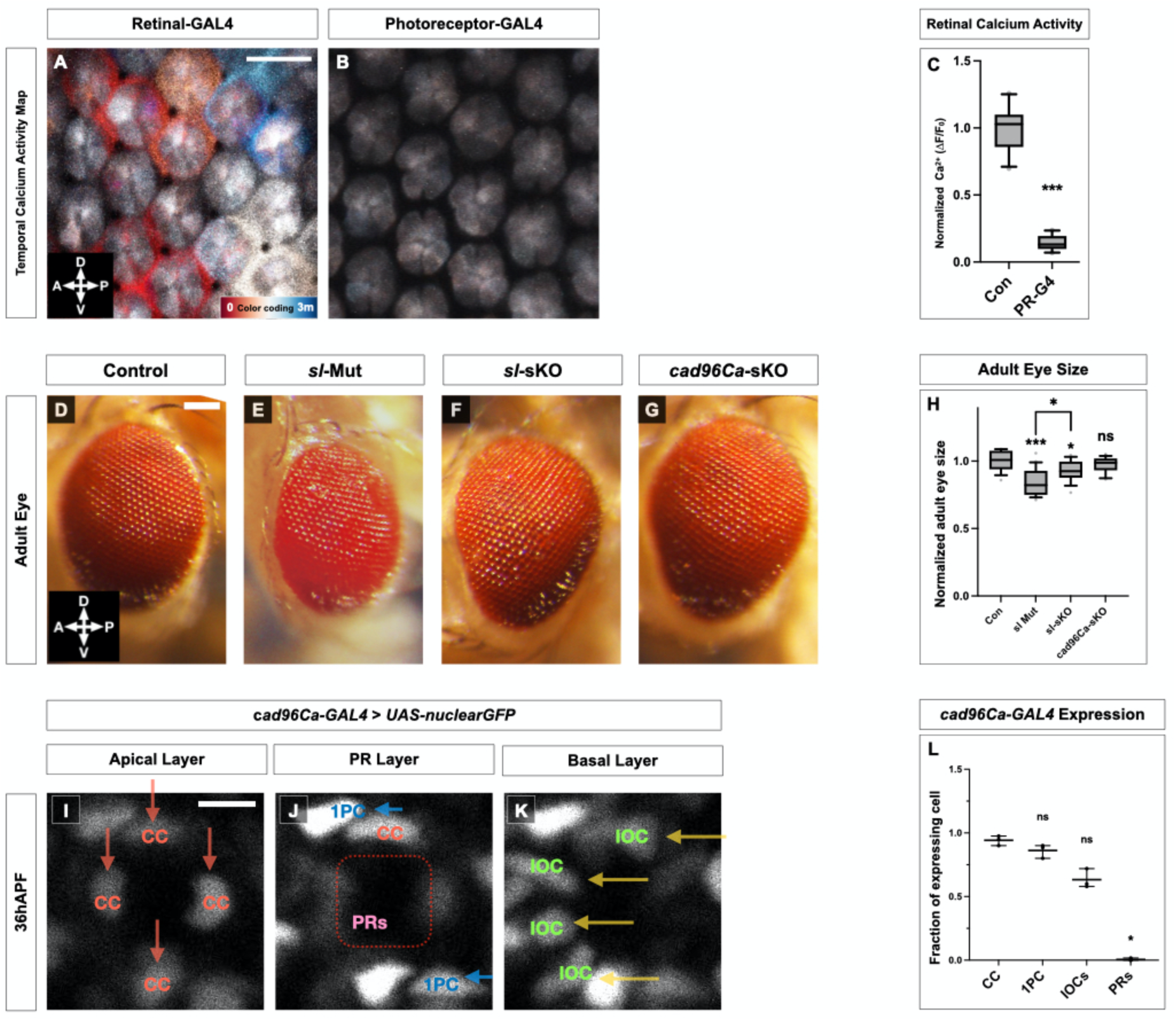
**S3A, B.** Representative images of retinal wave activity, displayed as color-coded 3-minute temporal calcium activity maps from pan-retinal GAL4 (longGMR-GAL4) **(A)** and a photoreceptor-specific GAL4 (Chaoptin-GAL4) **(B)** retinas during stage 2 (**A**) or corresponding time window (**B**). **B** shows only basal calcium signal without meaningful activity. Scale bar: 10 μm **(A, B).** See also **Movie S2**. **S3C.** Quantification of retinal calcium activity from **S3A, B**. n ≥ 8 retinas, ***: p < 0.001, error bars: SD. **S3D-G.** Representative images of adult eyes from control **(D),** *sl* null mutant **(E),** *sl*-*sKO* **(F),** and *cad96Ca*-*sKO* **(G)**. **S3H.** Quantification of adult eye size from **SD-G.** n ≥ 14 retinas, ns: p≥ 0.05, *: p < 0.05, ***: p < 0.001, error bars: SD. **S3I-K.** Live imaging snapshots of retinas at 36hAPF, using a nuclear GFP reporter driven by *cad96Ca-GAL4*. Each panel represents a single Z-section of retinal layers from apical **(I),** photoreceptor **(J),** and basal **(K)** layers. CC = cone cell, PR = photoreceptor, 1PC = primary pigment cell, IOC = interommatidial cell. Scale bar: 10 μm **(I-K)**. **S3L.** Quantification of the fraction of *cad96Ca-GAL4* expressing cell-types from **S3I-K.** n ≥ 93 cells from 3 retinas, ns: p≥ 0.05, *: p < 0.05.

**Fig. S4 (A–Z) related to Fig. 4.**
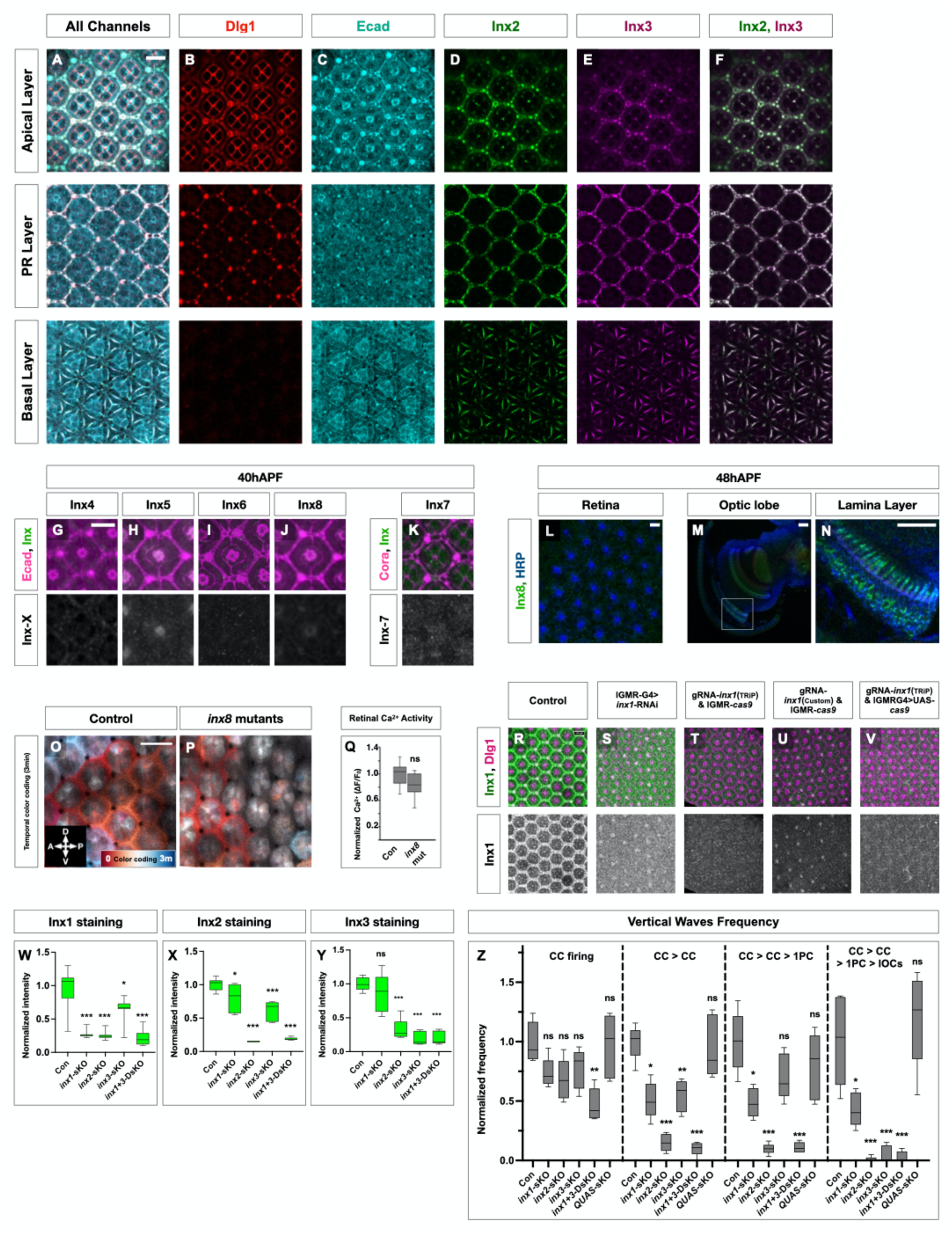
**S4A-F.** Representative immunostaining images of Inx2 and Inx3 in 40hAPF retinas. These Z-stack images display the apical layer **(A-F tops)**, photoreceptor (PR) layer **(A-F middles)**, and basal layer **(A-F bottoms)**. Gap junction plaques containing Inx2 and Inx3 are observed at retinal support cell junctions, labeled with either Dlg1 (**red**) or E-cadherin (**cyan**). No Inx2 or Inx3 plaques were detected around PR cell bodies, indicated by empty space in the center of the ommatidium **(F middle)**, confirming that gap junctions are primarily in retinal non-neuronal cells. Scale bar: 10 μm **(A-F)**. **S4G-K.** Representative immunostaining images of Inx4–8 (Inx-X) in 40hAPF retinas, with innexins **(green for tops, white for bottoms)** and adherens junctional marker E-cadherin **(magenta for G-J tops)** or septate junctional marker Coracle **(magenta for K top)**. No significant Inx 4–8 signals were detected at ommatidial junctions in 40hAPF retina, except weak Inx5 signal from early rhabdomeres **(H)**. Scale bar: 10 μm **(G-K)**. **S4L-N.** Immunostaining of Inx8 in retinas **(L)** and photoreceptor axonal terminals **(M, N)** at 48hAPF, using Inx8 **(green)** and neuronal membrane marker HRP **(blue)**, respectively. No meaningful signal was detected in the retina **(L)**, suggesting the subcellular localization of Inx8 in PRs is confined to axonal terminals located in the optic lobe **(M, N)**. Scale bar: 10 μm **(L-N). S4O, P.** Representative images of retinal wave activity, displayed as color-coded 3-minute temporal calcium activity maps from control **(O)** and *inx8* mutant retinas **(P)** during stage 2 waves. Scale bar: 10 μm **(O, P).** See also **Movie S3**. **S4Q.** Quantification of retinal calcium activity from **S4O, P.** n ≥ 10 retinas, ns: p ≥ 0.05, error bars: SD. **S4R-V.** Representative retinal images of transgenic *inx1* mutants at 40hAPF, including control **(R),** *inx1-RNAi* **(S),** *gRNA-inx1* (BDSC version) with *longGMR-Cas9* **(T),** *gRNA-inx1* (custom version) with *longGMR-Cas9* (referred to as ***inx1*-*sKO***) **(U),** and *gRNA-inx1* (BDSC version) with *longGMR-GAL4 > UAS-Cas9* **(V)**, using Inx1 **(green)** and junctional marker Dlg1 **(magenta)**, respectively. The images are Z-stacks of the retinal apical layer. Scale bar: 10 μm **(R-V)**. **S4W-Y.** Quantification of Inx1 **(W),** Inx2 **(X),** and Inx3 **(Y)** immunostaining signals in *inx*s-*sKO* mutant retinas from Fig. 4E**-I**. n ≥ 6 retinas, ns: p ≥ 0.05, *: p < 0.05, **: p < 0.01, ***: p < 0.001, error bars: SD. **S4Z.** Quantification of vertical pathway frequency for each step indicated above in *inx* mutant retinas during late stage 1. All values were normalized to control. n ≥ 200 ommatidia from 5 retinas, ns: p ≥ 0.05, *: p < 0.05, **: p < 0.01, ***: p < 0.001, error bars: SD.

**Fig. S5 (A–P) related to Fig. 5.**
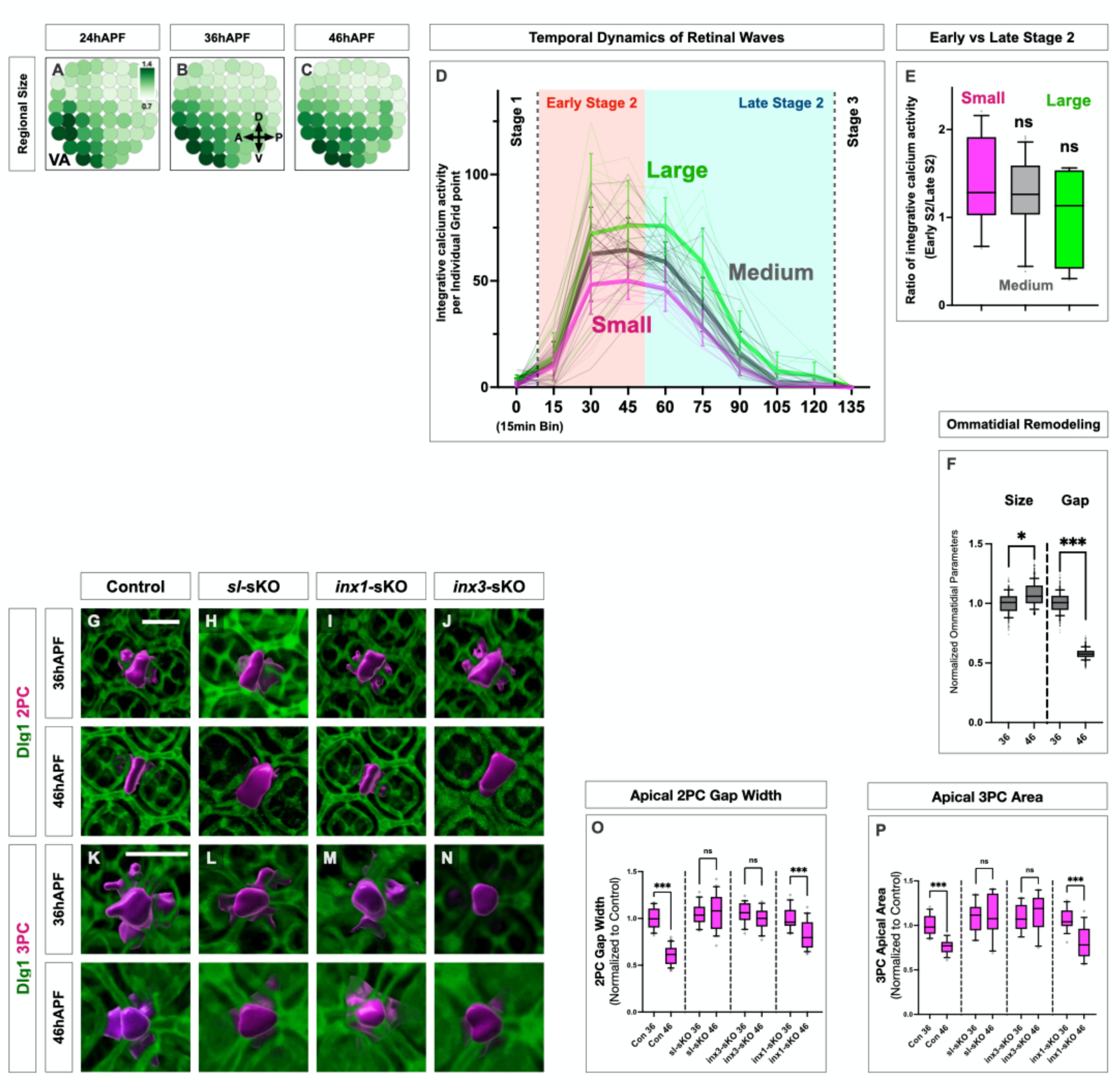
**S5A-C.** Ommatidial configuration grid of size distribution in control retinas at 24hAPF (**A**), 36hAPF (**B**), and 46hAPF (**C**). Each circle represents a group of ommatidia in an 8×8 grid. Color indicates average ommatidial size per grid point, showing a strong ventral-anterior bias at mid-pupal stages. n ≥ 3 control retinas. **S5D.** Trace of calcium activity during retinal waves. Each time point represents integrative calcium activity over 15-min intervals. Thin lines indicate activity for each 8×8 grid point, while thick lines show average activity for grid groups with large (80–100%, green), small (0–20%, magenta), and medium (20–80%, gray) ommatidial sizes. n ≥ 11 retinas, error bars: SD. **S5E.** Quantification of integrative calcium activity between early and late phases of stage 2 from S5B. Y-axis shows the normalized ratio of early to late stage 2 activity, indicating no significant differences between size groups. n ≥ 11 retinas; ns: p ≥ 0.05; error bars: SD. **S5F.** Quantification of individual ommatidial size and inter-ommatidial gaps between 36hAPF (blue) and 46hAPF (red) from ommatidia labeled with Dlg1 (Fig. 5O). Definition of each parameter is shown in Fig. 5M, N. n ≥ 288 ommatidia from n ≥ 9 retinas, *: p < 0.05, ***: p < 0.001, error bars: SD. **S5G-N.** Representative 3D-reconstructed images of secondary and tertiary pigment cells (2PCs, 3PCs) at different stages **(36hAPF: G-N tops; 46hAPF: G-N bottoms)** in different genetic backgrounds, labeled with membrane-tethered MCFO-epitope tag (magenta) and Dlg1 (green). Scale bar: 10 μm **(G-N)**. **S5O, P.** Quantification of apical contraction of gap-width in 2PCs as related to control size **(O)** and 3PC area **(P).** n ≥ 21 2PCs, n ≥ 11 3PCs from ≥10 retinas per genotype, *: p < 0.05, **: p < 0.01, ***: p < 0.001, error bars: SD.

**Fig. S6 (A–E) related to Fig. 5-7.**
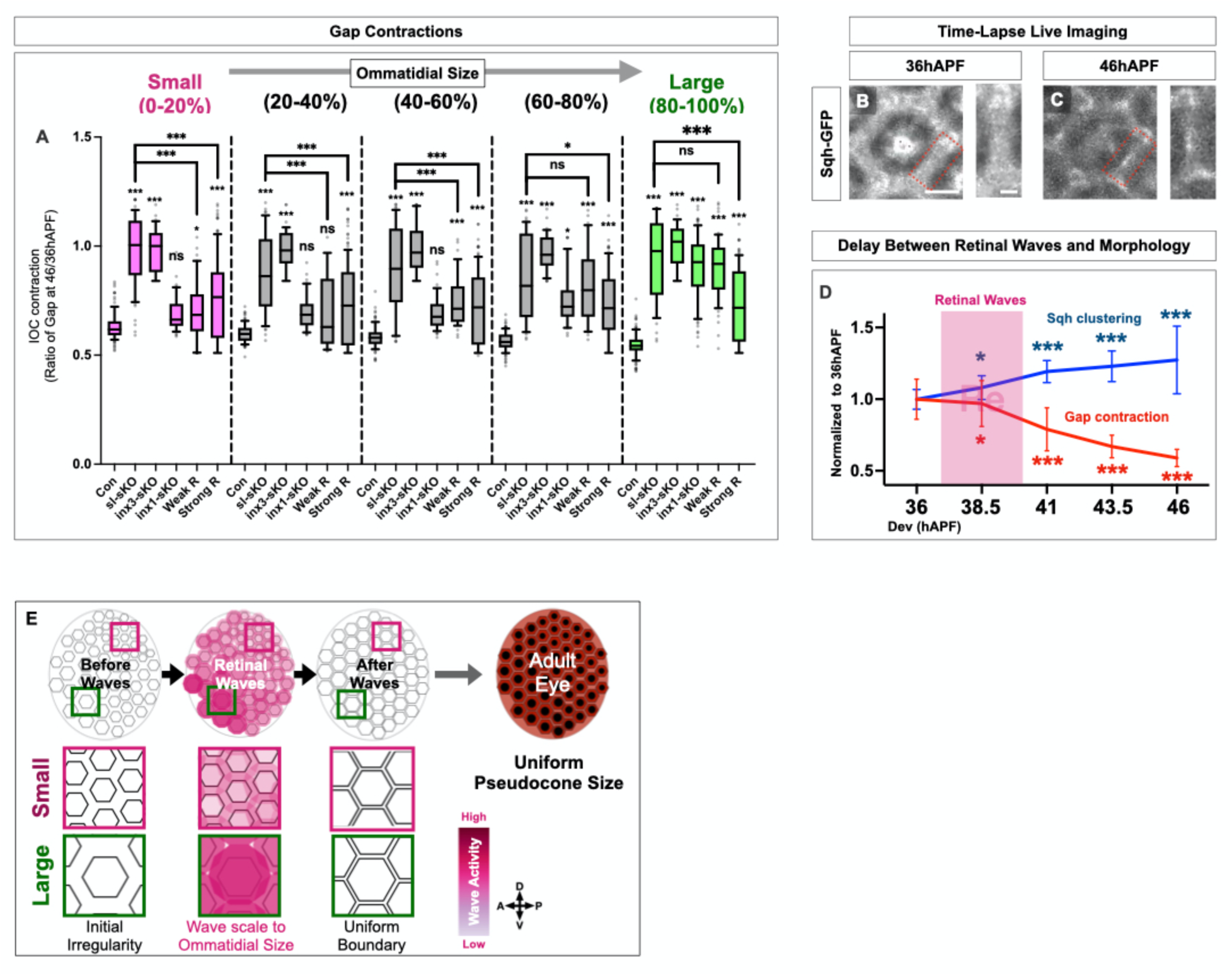
**S6A.** Quantification of inter-ommatidial gap contraction with various sizes of ommatidia using Dlg1 immunostaining (Fig. 5F**-L**). The Y-axis represents IOC contraction, calculated as the ratio of 46hAPF inter-ommatidial gap size to that of 36hAPF for each group. n ≥ 26 ommatidia from n ≥ 16 retinas per size group, ns: p ≥ 0.05, *: p < 0.05, **: p < 0.01, ***: p < 0.001, error bars: SD. **S6B, C.** Time-lapse images of a single ommatidium at 36hAPF (**B**) and 46hAPF (**C**) in the control retina, showing Sqh-GFP puncta in the apical side of inter-ommatidial junction. The red box and side insets highlight apical interommatidial junctions with Sqh-GFP clusters, indicating actomyosin network formation during retinal waves. **S6D**. Overlay of boundary contraction, Myosin II clustering, and retinal wave periods Traces show boundary contraction (apical Dlg1, red) and Sqh-GFP clustering (blue) across developmental stages, with the retinal wave period highlighted in magenta. Values are normalized to the average at 36hAPF; statistical comparisons are to 36hAPF. n ≥ 99 (boundary contraction) from 20 retinas; n ≥ 50 (Sqh-GFP clustering) from ≥5 retinas. ns: p ≥ 0.05; *p < 0.05; ***p < 0.001, error bars: SD. **S6E.** Summary diagram illustrating how retinal waves coordinate uniform pseudocone formation in the adult eye through apical remodeling scaled by ommatidial size. At 36hAPF, the retina exhibits regional ommatidial size variation, accompanied by interommatidial gap differences. Retinal waves promote regional calcium signaling proportionate to ommatidial size, which induces dose-dependent interommatidial gap contraction. This compensating process ensures the formation of uniform ommatidial boundaries by 46hAPF, which are essential for uniform pseudocone development in the adult retina.

**Table S1.**

| Gene Symbol | Gene Name | FlyBase ID | Mammalian Homolog | Genetic Manipulation | Gross Retinal Morphology | Early Retinal Waves |
| --- | --- | --- | --- | --- | --- | --- |
| **alk** | Anaplastic lymphoma kinase | FBgn0040505 | ALK | UAS-Dominant negative (BL#92968) | normal | normal |
| **cad96Ca** | Cad96Ca/Stitcher | FBgn0022800 | Not known | guide RNA (Customized version) | normal | Severe disruption |
|  |  |  |  | UAS-RNAi (BL#55877) | normal | Mild disruption |
| **ddr** | Discoidin domain receptor | FBgn0053531 | DDR1 and DDR2 | UAS-RNAi (BL#55906) | normal | normal |
| **dnt** | Doughnut on 2 | FBgn0024245 | RYK | guide RNA (Customized version) | normal | normal |
|  |  |  |  | UAS-RNAi (BL#67251) | normal | normal |
| **drl** | Derailed | FBgn0015380 | RYK | guide RNA (Customized version) | normal | normal |
|  |  |  |  | UAS-RNAi (BL#39002) | normal | normal |
| **egfr** | breathless | FBgn0003731 | EGFR | UAS-Dominant negative (BL#5364) | Severe disruption | Mild disruption |
| **eph** | Eph receptor tyrosine kinase | FBgn0025936 | EPHA and EPHB | guide RNA (Customized version) | normal | normal |
|  |  |  |  | UAS-RNAi (BL#60006) | normal | normal |
| **inR** | Insulin-like receptor | FBgn0013984 | INSR/IGF1R | UAS-Dominant negative (BL#8253) | normal | normal |
| **nrk** | Neurospecific receptor kinase | FBgn0020391 | MuSK | UAS-RNAi (BL#56936) | normal | normal |
| **otk** | off-track | FBgn0004839 | Wnt4 | guide RNA (Customized version) | normal | normal |
| **pvr** | PDGF- and VEGF-receptor related | FBgn0032006 | VEGFR and PDGFR | guide RNA (Customized version) | Mild disruption | Mild disruption |
|  |  |  |  | UAS-Dominant negative (BL#58431) | Mild disruption | Mild disruption |
| **ret** | Ret oncogene | FBgn0011829 | RET | UAS-RNAi (BL#55872) | normal | normal |
| **sev** | sevenless | FBgn0003366 | Not known | UAS-RNAi (BL#55866) | normal | normal |
| **tie** | Tie-like receptor tyrosine kinase | FBgn0014073 | Not known | guide RNA (Customized version) | normal | normal |
| **tor** | torso | FBgn0003733 | Not known | guide RNA (Customized version) | normal | Mild disruption |
|  |  |  |  | UAS-RNAi (BL#58312) | normal | normal |
| **Control** | QUAS sequence |  |  | guide RNA (Customized version) | normal | normal |

## Notes

### Competing Interest Statement

The authors have declared no competing interest.

### Summary of Updates

Revised supplementary control results and improved figure labeling, including color-blind-friendly adjustments.

